# Shared early impairments of medial entorhinal cortex function across distinct Alzheimer’s disease etiologies

**DOI:** 10.64898/2026.05.09.723018

**Authors:** Beate Throm, Elke Fuchs, Jing-Jie Peng, Hannah Ostermann, Wolfgang Wick, Kevin Allen, Hannah Monyer

**Affiliations:** Medical Faculty of Heidelberg University and German Cancer Research Center (DKFZ); Heidelberg, Im Neuenheimer Feld 280, 69120, Germany; Neurology Clinic of the University Hospital Heidelberg, Im Neuenheimer Feld 400, 69120 Heidelberg

**Keywords:** Grid cells, head-direction cells, theta oscillations, medial septum, in vivo electrophysiology, object-vector coding, spatial memory, 5xFAD, Tau P301S

## Abstract

The medial entorhinal cortex (MEC) harbors diverse spatially and object-tuned cell types essential for spatial coding and episodic memory, and is among the earliest regions affected in neurodegenerative diseases. Here, we examined how early neurodegeneration alters MEC cellular coding, network dynamics, and behavior in two Alzheimer mouse models of distinct etiology. Specifically, we performed in vivo electrophysiology, immunohistochemistry and behavioral assays in models that reflect the manifestation of a tauopathy and amyloidopathy, respectively, to identify common impairments. In both models, the earliest deficits manifested as instability of spatial context representations at the cellular and behavioral level. These impairments preceded model-specific disruptions in grid cell coding and altered theta oscillations. We further identified reduced parvalbumin-positive (PV^+^) septal projections as a likely contributor to MEC dysfunction. In contrast, object-vector coding remained intact, highlighting spatial context instability as an early marker of MEC impairment.

## Introduction

The medial entorhinal cortex (MEC), located at the interface between neocortical areas and hippocampus, serves as a hub for spatial coding and episodic memory. This is reflected by the presence of diverse spatially-tuned cell types such as grid cells ^1^, head-direction (HD) cells ^2^ and border cells ^3^, which together generate a stable, yet flexible representation of space across different spatial contexts ^4–6^. Within a given environment, grid cells encode a self-motion updated metric of space through their periodic hexagonal firing pattern, while HD cells provide a stable directional reference and border cells anchor spatial representations to environmental boundaries. Major changes in an environment explored by an animal, lead to a coherent rotation in the grid cell network ^7^, accompanied by corresponding rotations in the preferred firing direction of HD cells and shifts in the firing fields of border cells ^2,3,8^. This coordinated interplay among spatially-tuned cell types in the MEC ensures the continuity and adaptability of spatial representations across spatial contexts, supporting navigation and episodic memory ^9^.

The temporal organization of spatial coding in the MEC is provided by theta oscillations, a prominent 6–12 Hz rhythm characteristic of the rodent hippocampal–entorhinal system ^10^. Thus, the ordered firing of spatially-tuned cells with respect to the theta cycle, not only in the MEC but in the entire hippocampal formation, is one of the best predictors of the animal’s present and upcoming location ^11–15^. Importantly, an analogous theta-band oscillatory pattern was found in intracranial recordings of the human medial entorhinal cortex during spatial navigation and memory tasks ^16–19^. Converging histological and electrophysiological studies provided strong evidence that the medial septum (MS) subserves as pacemaker of hippocampal–entorhinal theta activity via its rhythmic inhibitory and cholinergic input to the MEC ^20–23^. Notably, optogenetic activation of parvalbumin-positive (PV⁺) GABAergic septal projections suffices to entrain theta-frequency oscillations in MEC and hippocampus ^24,25^. Accordingly, chemogenetic silencing of septal PV⁺ input to MEC disrupted endogenous theta oscillations and impaired reference memory performance, underscoring the functional importance of septal-driven theta rhythms for memory-guided behavior ^26^.

Recent studies have added to the list of spatially-tuned cells in MEC yet other cell types that are object-tuned: Object-vector cells fire at a certain distance and orientation from objects ^27^, and object-tuned fast-spiking cells are fast-spiking neurons with enhanced firing activity around objects ^28^. This vectorial-based representation of space could be used in landmark-based navigation ^29^.

At the behavioral level, the significance of MEC for spatial coding and episodic memory has been highlighted in numerous studies involving lesions or (opto)genetic manipulations ^30–36^. For instance, lesioning MEC led to selective deficits in an object-displacement test, but not in an object-identification task ^31^.

In neurodegenerative diseases such as Alzheimer’s disease (AD), the MEC is amongst the first brain regions affected, accounting for the earliest cognitive deficits such as spatial navigation and episodic memory deficits ^37–41^. In particular, a path integration test proved to have a high sensitivity and specificity in distinguishing between healthy controls and mild cognitive impairment, a clinical condition preceding dementia ^42^. The characterization of different mouse models, including mice with diverse amyloid and tau pathologies, revealed quite distinct impairments at the cellular level and at the network level in vivo in MEC ^43–50^. While grid cell disruption seems to emerge as a common feature at least in advanced stages of the pathology ^47–50^, it remains unclear whether and how the complex interplay of spatially-tuned cells across spatial contexts and object-vector coding is affected at early stages of neurodegeneration.

Here we used two mouse models of neurodegeneration to probe for common signs of early MEC malfunction at the functional and behavioral level. Specifically, we used Tau P301S and 5xFAD mice as a model reflecting a tauopathy and an amyloid pathology respectively. By combining in vivo electrophysiological recordings with immunohistochemistry and behavioral assays, we sought to identify common features in the two mouse models of AD at the cellular and network level. Furthermore, we tested how these translate into a behavioral deficit that may serve as an early marker at incipient stages of a neurodegenerative process affecting the MEC.

## Results

### Grid cell malfunction in mature amyloid and tau mice

To explore the impact of tau and amyloid pathology on different cell types in MEC in the incipient phase of neurodegeneration, we performed electrophysiological experiments with 6 different cohorts, comprising wild type (“control”), Tau P301S (“tau”) and 5xFAD mice (“amyloid”) at 5-6 months of age (“young” cohorts) as well as 8-9 months of age (“mature” cohorts). We deliberately chose age-ranges before neuronal loss in the MEC had occurred (**Supp. Fig. 1**). For this purpose, we implanted a total of 61 mice with silicon probes targeted at the superficial layers of MEC (young control: 13 mice, 133 sessions; young tau: 11 mice, 97 sessions; young amyloid: 7 mice, 62 sessions; mature control: 11 mice, 73 sessions; mature tau: 8 mice, 80 sessions; mature amyloid: 11 mice, 99 sessions; **Supp. Tables 1.1-1.6**). Histology confirmed that the vast majority of probe shanks in all 6 cohorts were found in the superficial layers of MEC (**Supp. Fig. 2 a-b**).

**Fig. 1:**
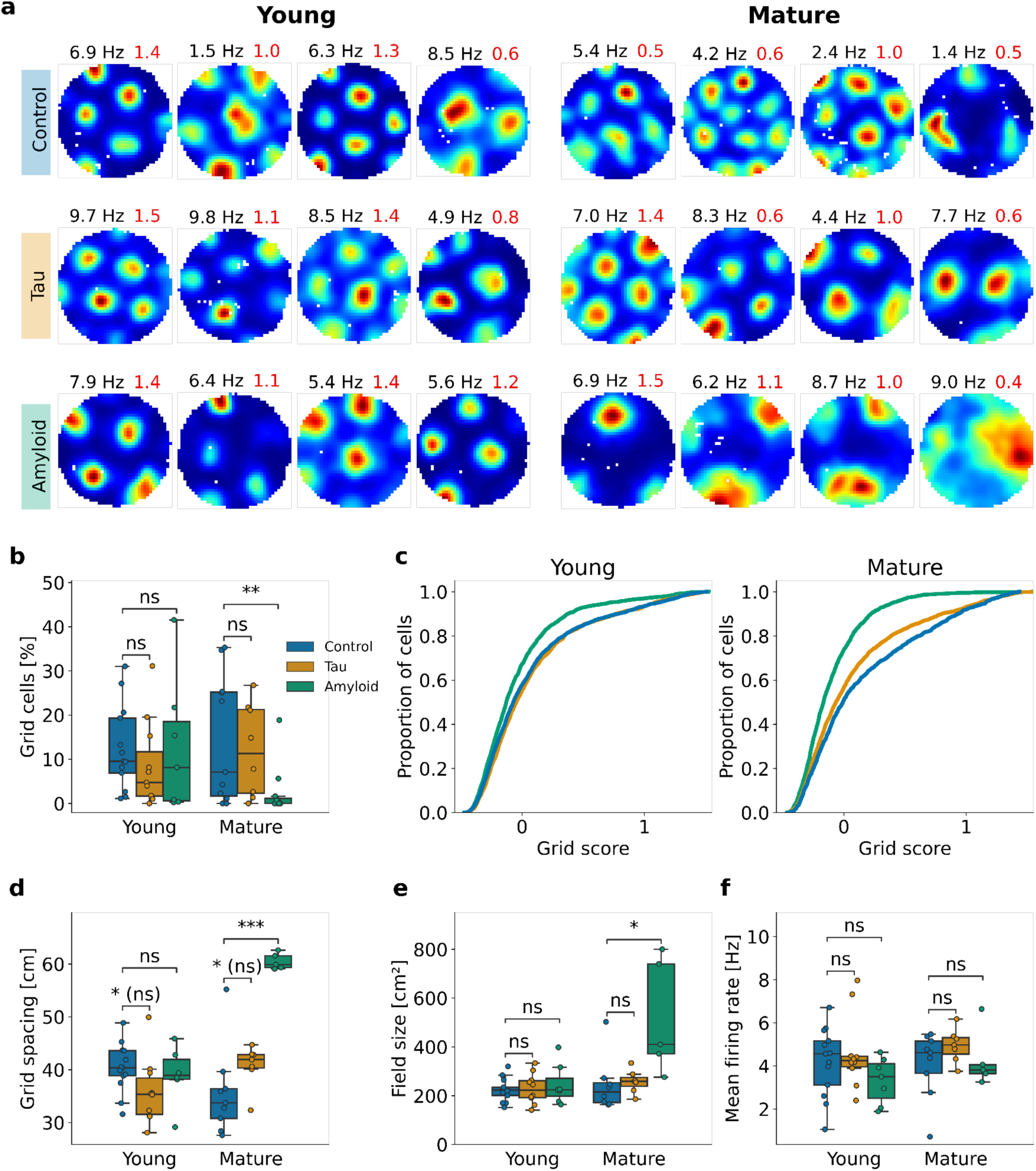
Reduction in grid cell number and larger grid scale in mature amyloid mice. (a) Example grid cells in control, tau and amyloid mice. 4 representative examples are shown for each group. Peak firing rate (left) and grid score (right, red) are shown above each firing rate map. (b) Percentage of grid cells for each mouse. Each mouse is represented by a dot. The legend applies to (b)-(f). Mann-Whitney-Wilcoxon test, one-sided with Holm-Bonferroni correction, statistical unit: mouse: young control vs. young tau: p = 0.11, U = 93; young control vs. young amyloid: p = 0.27, U = 54; mature control vs. mature tau: p = 0.42, U = 47; mature control vs. mature amyloid: p = 0.0077, U = 97 (c) Distribution of grid scores among all non-fast-spiking cells, plotted separately for young and mature cohorts. Mann-Whitney-Wilcoxon test, one-sided with Holm-Bonferroni correction, statistical unit: cell: young control (n = 3045) vs. young tau (n = 1778): p = 0.99, U = 2605,280; young control vs. young amyloid (n = 2154) : p = 3.60*10^-14^, U = 3678,470; mature control (n = 1326) vs. mature tau (n = 2893): p = 1.04*10^-6^, U = 2092,327; mature control vs. mature amyloid (n = 1830) : p = 2.6*10^-52^, U = 1596,704 (d-f) Boxplots showing the grid spacing, mean field size and mean firing rate of grid cells. Each data point is the median of the grid cells recorded in one mouse. (d) Grid spacing: Mann-Whitney-Wilcoxon test, two-sided with Holm-Bonferroni correction, statistical unit: mouse: young control vs. young tau: p = 0.038 (n.s. after Holm-Bonferroni correction), U = 99; young control vs. young amyloid: p = 0.58, U = 46; mature control vs. mature tau: p = 0.042 (n.s. after Holm-Bonferroni correction), U = 12; mature control vs. mature amyloid: p = 0.0010, U = 0 (e) Mean field size: Mann-Whitney-Wilcoxon test, two-sided with Holm-Bonferroni correction, statistical unit: mouse: young control vs. young tau: p = 1.0, U = 65; young control vs. young amyloid: p = 0.84, U = 42.5; mature control vs. mature tau: p = 0.23, U = 17; mature control vs. mature amyloid: p = 0.011, U = 30 (f) Mean firing rate: Mann-Whitney-Wilcoxon test, two-sided with Holm-Bonferroni correction, statistical unit: mouse: young control vs. young tau: p = 0.98, U = 66; young control vs. young amyloid: p = 0.16, U = 64; mature control vs. mature tau: p = 0.30, U = 21; mature control vs. mature amyloid: p = 0.70, U = 26

**Fig 2:**
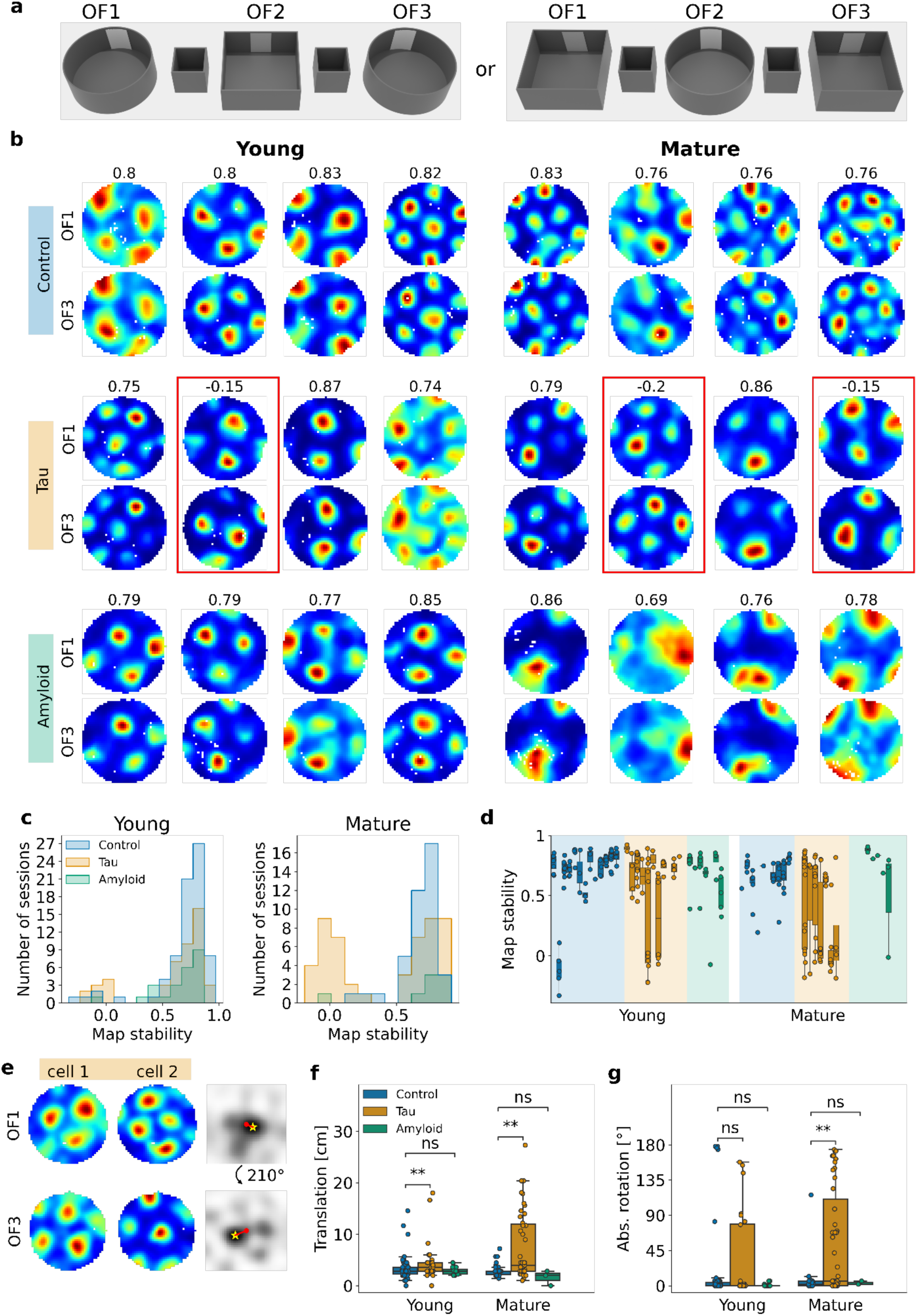
Inter-trial map stability of grid cells is reduced in mature tau mice. (a) Schematic of the A-B-A protocol. Mice were randomly assigned to start in the circular box or the square box. The first open-field trial (OF1) was followed by a rest box trial before the circular box was exchanged for the square box and vice versa (OF2). After another rest box trial, the session was concluded by an open-field trial in the start box (OF3). (b) Firing rate maps of example grid cells in the first and last open field trial (OF1 and OF3). The Pearson correlation coefficient between the firing rate maps of the two trials is shown above the maps of each neuron. Red boxes mark examples with a map cross-correlation < 0.4. (c) Histograms of inter-trial map stability for control, tau and amyloid mice. Data from young and mature mice are plotted separately. Each data point is the median Pearson correlation coefficient between maps of grid cells recorded in one session. The legend applies to (b)-(e). Mann-Whitney-Wilcoxon test, one-sided, statistical unit: session: young control vs. young tau: p = 0.065, U = 2106; young control vs. young amyloid: p = 0.095, U = 1047; mature control vs. mature tau: p = 0.0074, U = 1325; mature control vs. mature amyloid: p = 0.97, U = 93 (d) Boxplot showing the inter-trial map stability. Each bar shows map stability for one mouse, with each data point representing the median stability of all recorded grid cells in one session. Fewer data points are available in mature amyloid mice because of the small number of sessions with grid cells. (e) Schematic explaining how grid cell rotation was calculated: Left: Firing rate maps of the 2 grid cells for the first and last open field trial (OF1, OF3). Right: cross-correlograms of the respective firing rate maps for OF1 (upper row) and OF3 (lower row). For better orientation, the center and the closest peak are marked with a red dot and a yellow asterisk respectively.; the red line represents the offset vector. The resulting rotations in a counter-clockwise direction are printed between the cross-correlograms. (f) Grid cell phase shift between the first and the last open field trial. Each dot represents a session with at least 2 co-recorded grid cells. Mann-Whitney-Wilcoxon test, one-sided, statistical unit: session: young control vs. young tau: p = 0.01, U = 530; young control vs. young amyloid: p = 0.62, U = 493; mature control vs. mature tau: p = 0.0012, U = 388.5; mature control vs. mature amyloid: p = 0.11, U = 74.5 (g) Absolute grid cell rotation between the first and the last open field trial (up to 180° in either direction). Each dot represents a session with at least 2 co-recorded grid cells. Mann-Whitney-Wilcoxon test, one-sided, statistical unit: session: young control vs. young tau: p = 0.71, U = 762; young control vs. young amyloid: p = 0.067, U = 578; mature control vs. mature tau: p = 0.0032, U = 420; mature control vs. mature amyloid: p = 0.64, U = 40

We first tested grid cell integrity in a familiar environment using a simple open-field paradigm. Grid cells in mature amyloid mice were almost entirely absent and the few remaining ones were altered compared to those in the 5 other cohorts (**Fig. 1 a**). Indeed, the proportion of grid cells in mature amyloid mice was significantly reduced compared to mature control mice (**Fig. 1 b**). When analyzing the distributions of grid scores among all non-fast-spiking cells, we found a marked reduction in grid scores already in young amyloid mice and also in mature tau mice (**Fig. 1 c**). In addition, the remaining grid cells recorded in mature amyloid mice displayed a greater grid spacing and field size (**Fig. 1 d-e**). The recording depth (distance of the shank tip from the dorsal border of MEC in the dorsoventral axis) did not differ across cohorts (**Supp. Fig. 2 c-f**). Other basic properties such as the mean firing rate (**Fig. 1 f**), the information score and the peak firing rate (data not shown) were not altered. To further test the functional properties of grid cells, we analyzed their relation to theta and phase precession. Interestingly, grid cells in both mature tau and amyloid mice showed increased theta modulation, while only mature amyloid mice exhibited reduced phase precession slopes (**Supp. Fig. 3**). In sum, at around 8-9 months of age, grid cells in mice with an amyloid pathology exhibited quantitative and qualitative alterations: Thus, grid cell number per se was reduced and the remaining grid cells exhibited large firing fields and altered phase precession. In contrast, grid cells in tau mice remained largely intact throughout the observation time.

**Fig. 3:**
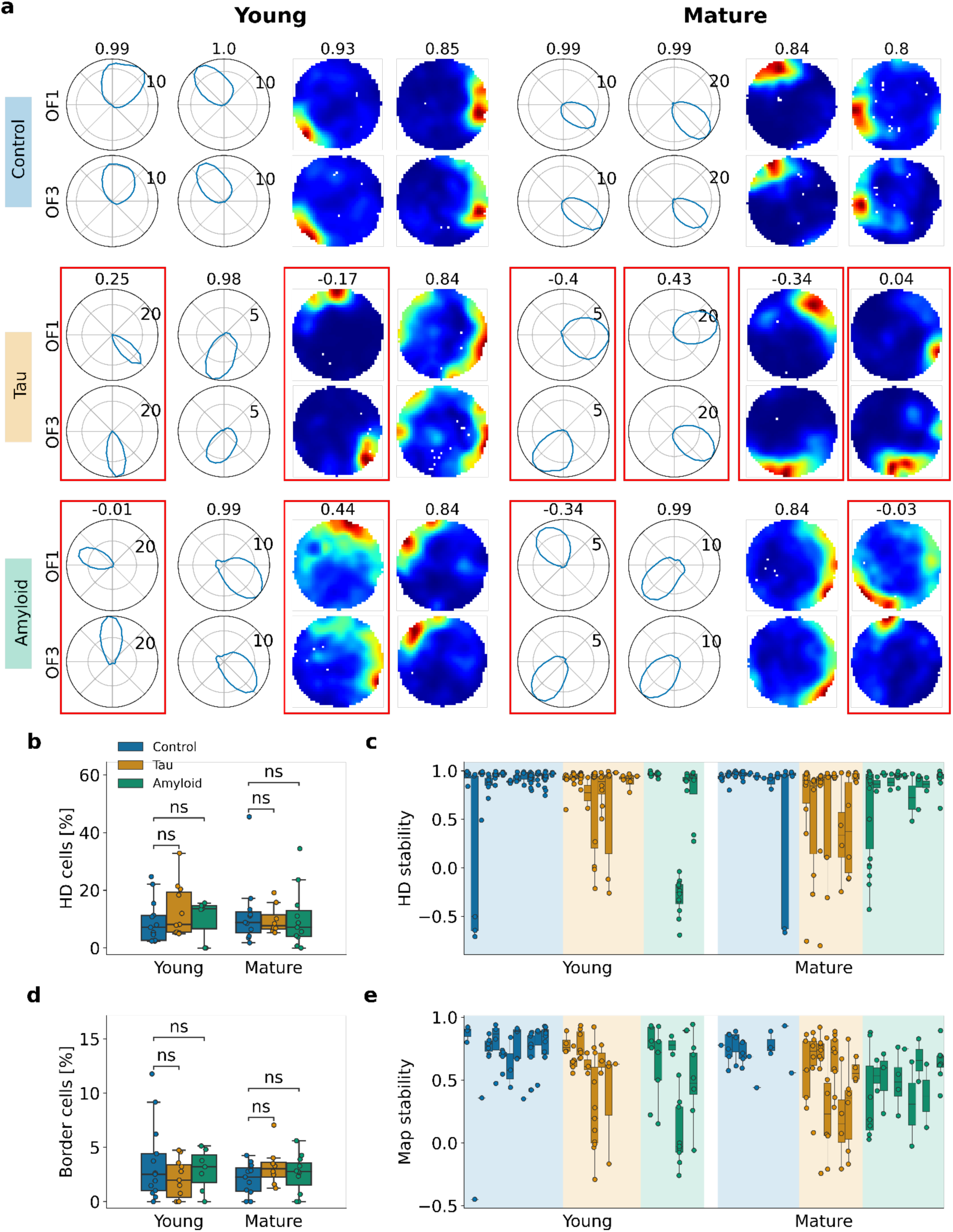
Head-direction and border cells in tau and amyloid mice exhibit reduced inter-trial stability already in young AD mice. (a) Examples of HD cells (polar HD histograms) and border cells (firing rate maps) in the first and last open field trial (OF1 and OF3) for each cohort. The Pearson correlation coefficient between firing rate maps or HD rate histograms is shown above map pairs. Red boxes mark examples with a correlation coefficient < 0.5. (b) Percentage of HD cells for each mouse. Each mouse is represented by a dot. The legend applies to (b)-(e). Mann-Whitney-Wilcoxon test, two-sided with Holm-Bonferroni correction, statistical unit: mouse: young control vs. young tau: p = 0.25, U = 51; young control vs. young amyloid: p = 0.69, U = 40; mature control vs. mature tau: p = 0.97, U = 45; mature control vs. mature amyloid: p = 0.60, U = 69 (c) HD stability of HD cells per mouse. Each bar represents the data from one mouse. Each dot is the median Pearson correlation coefficient of HD cells co-recorded in one session. Mann-Whitney-Wilcoxon test, one-sided with Holm-Bonferroni correction, statistical unit: session: young control vs. young tau: p = 0.080, U = 2945; young control vs. young amyloid: p = 0.14, U = 1908; mature control vs. mature tau: p = 0.012, U = 1571; mature control vs. mature amyloid: p = 0.030 (n.s. after Holm-Bonferroni correction), U = 1389 (d) Same as (b), but for border cells. Mann-Whitney-Wilcoxon test, two-sided with Holm-Bonferroni correction, statistical unit: mouse: young control vs. young tau: p = 0.47, U = 84.5; young control vs. young amyloid: p = 0.91, U = 43.5; mature control vs. mature tau: p = 0.26, U = 30; mature control vs. mature amyloid: p = 0.60, U = 52 (e) Same as (c), but for border cells. Mann-Whitney-Wilcoxon test, one-sided with Holm-Bonferroni correction, statistical unit: session: young control vs. young tau: p = 0.00076, U = 1392; young control vs. young amyloid: p = 0.0030, U = 1139; mature control vs. mature tau: p = 0.0054, U = 714; mature control vs. mature amyloid: p = 0.00071, U = 497

We next investigated the stability of the grid cell map using a circular arena and square box following an A-B-A-type protocol (**Fig. 2 a**). To counterbalance for potential environmental effects, mice of each cohort were assigned to two groups that were exposed to either the A-B-A or the B-A-B protocol. Young and mature control mice showed a high degree of map similarity (**Fig. 2 b**). However, the grid cells in some of the young tau mice and most of the mature tau mice repeatedly displayed a re-arrangement of the firing fields (**Fig. 2 b-d**).

Amyloid mice tended to have stable firing rate maps; yet in mature amyloid mice, the number of sessions with grid cells was much lower than in the other cohorts (8 sessions). When quantifying the inter-trial map stability, mature tau mice showed a significant reduction compared to mature control mice (**Fig. 2 c-d**). To investigate whether the re-arrangement of the grid pattern involved a rotation, a phase shift or both, we calculated the inter-trial grid rotation and translation for all sessions with at least 2 simultaneously recorded grid cells (**Fig. 2 e**, see Methods for details). Indeed, the reduced inter-trial stability in tau mice could be explained by a combination of a rotation and a translation of the grid cell network (**Fig. 2 f-g**). In a subset of sessions, we recorded several grid cells from 2 grid modules and we found similar rotations in both modules (**Supp. Fig. 4**). In sum, the rotation and the phase shift in the grid cell network in a subset of sessions from young and mature tau mice resulted in divergent representations of the same spatial context.

**Fig. 4:**
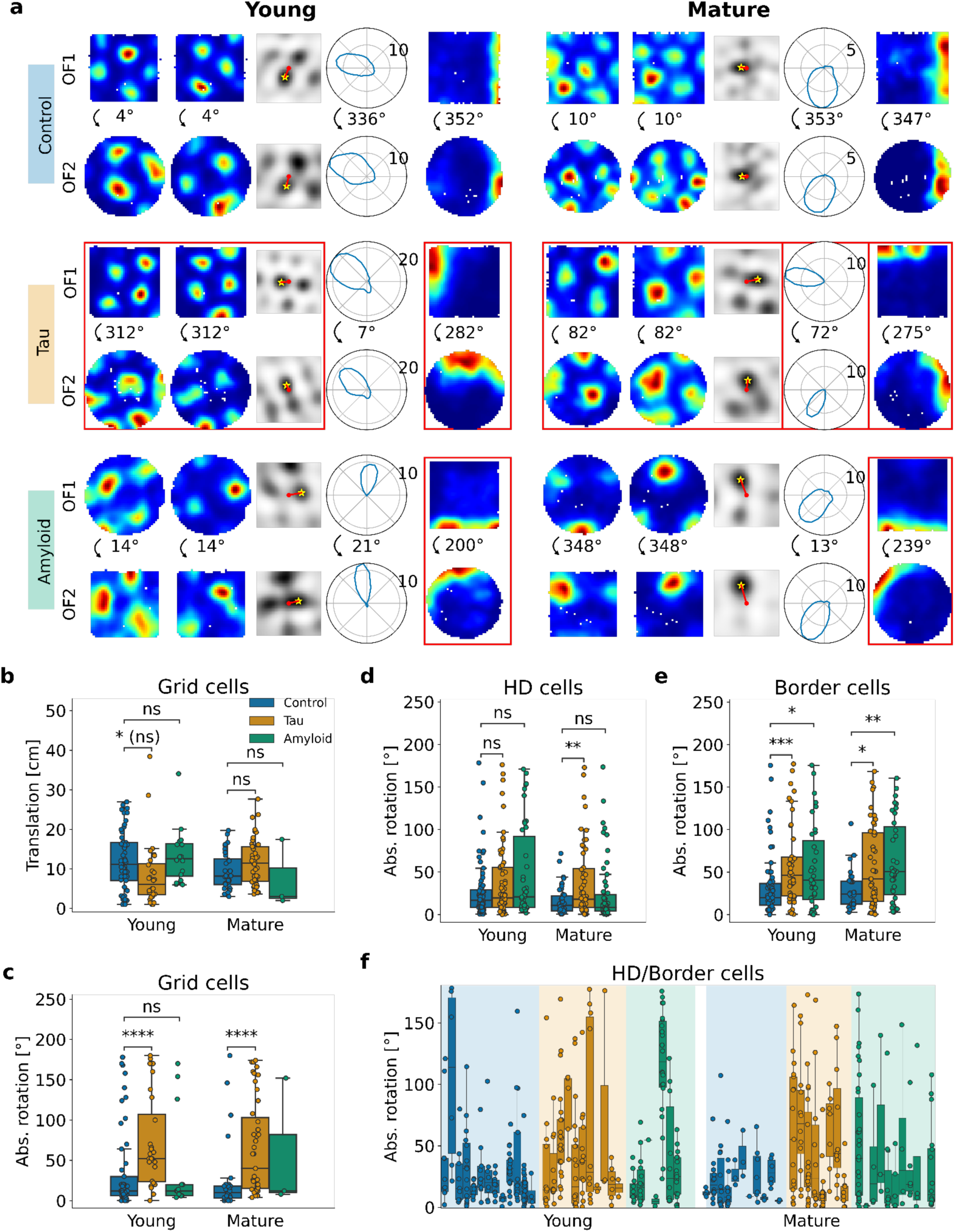
Grid cells, HD cells and border cells in tau mice and border cells in amyloid mice rotate between differently-shaped environments. (a) Firing rate maps of 2 example co-recorded grid cells and 1 example border cell as well as polar histograms of 1 example HD cell in the first and second open field trial (OF1 and OF2) for each cohort. The rotation of the cells (in counter-clockwise direction) is printed between the two trials. Red boxes mark examples with an absolute rotation of > 30°. (b) Translation of the grid pattern between the first and the second open-field trial for the young (left) and the mature cohort (right). Each dot is a session with at least 2 co-recorded grid cells. The legend applies to (b)-(e). Mann-Whitney-Wilcoxon test, two-sided with Holm-Bonferroni correction, statistical unit: session: young control vs. young tau: p = 0.019 (n.s. after Holm-Bonferroni correction), U = 1048; young control vs. young amyloid: p = 0.48, U = 402; mature control vs. mature tau: p = 0.11, U = 539; mature control vs. mature amyloid: p = 0.32, U = 65.5 (c) Absolute grid cell rotation between the first and the second open field trial (up to 180° in either direction). Each dot represents a session with at least 2 co-recorded grid cells. Mann-Whitney-Wilcoxon test, one-sided, statistical unit: session: young control vs. young tau: p = 9.6*10^-5^, U = 399; young control vs. young amyloid: p = 0.44, U = 443.5; mature control vs. mature tau: p = 2.4*10^-5^, U = 308.5 (d) HD rotation of HD cells. Each dot is the absolute circular median HD rotation of HD cells co-recorded in one session. Mann-Whitney-Wilcoxon test, one-sided with Holm-Bonferroni correction, statistical unit: session: young control vs. young tau: p = 0.13, U = 2308; young control vs. young amyloid: p = 0.073, U = 1428; mature control vs. mature tau: p = 0.0062, U = 811; mature control vs. mature amyloid: p = 0.66, U = 1188 (e) Same as (d), but for border cells. Mann-Whitney-Wilcoxon test, one-sided with Holm-Bonferroni correction, statistical unit: session: young control vs. young tau: p = 0.00024, U = 821; young control vs. young amyloid: p = 0.012, U = 853; mature control vs. mature tau: p = 0.016, U = 443; mature control vs. mature amyloid: p = 0.0035, U = 282 (f) Absolute rotation of HD and border cells per mouse for the 6 cohorts. Each bar represents the data from one mouse. Each dot is the absolute circular median rotation of HD cells and border cells recorded in one session.

### Functional impairment of HD and border cells in tau and amyloid mice

Next, we analyzed the integrity of other spatially-tuned and directionally-tuned cells. The proportion of HD cells and border cells as well as basic properties such as their peak firing rate and the HD or border score were similar to wild-type mice (**Fig. 3 a,b,d, Supp. Fig. 5**), suggesting that head-direction coding and border coding were preserved in young as well as mature tau and amyloid mice.

**Fig. 5:**
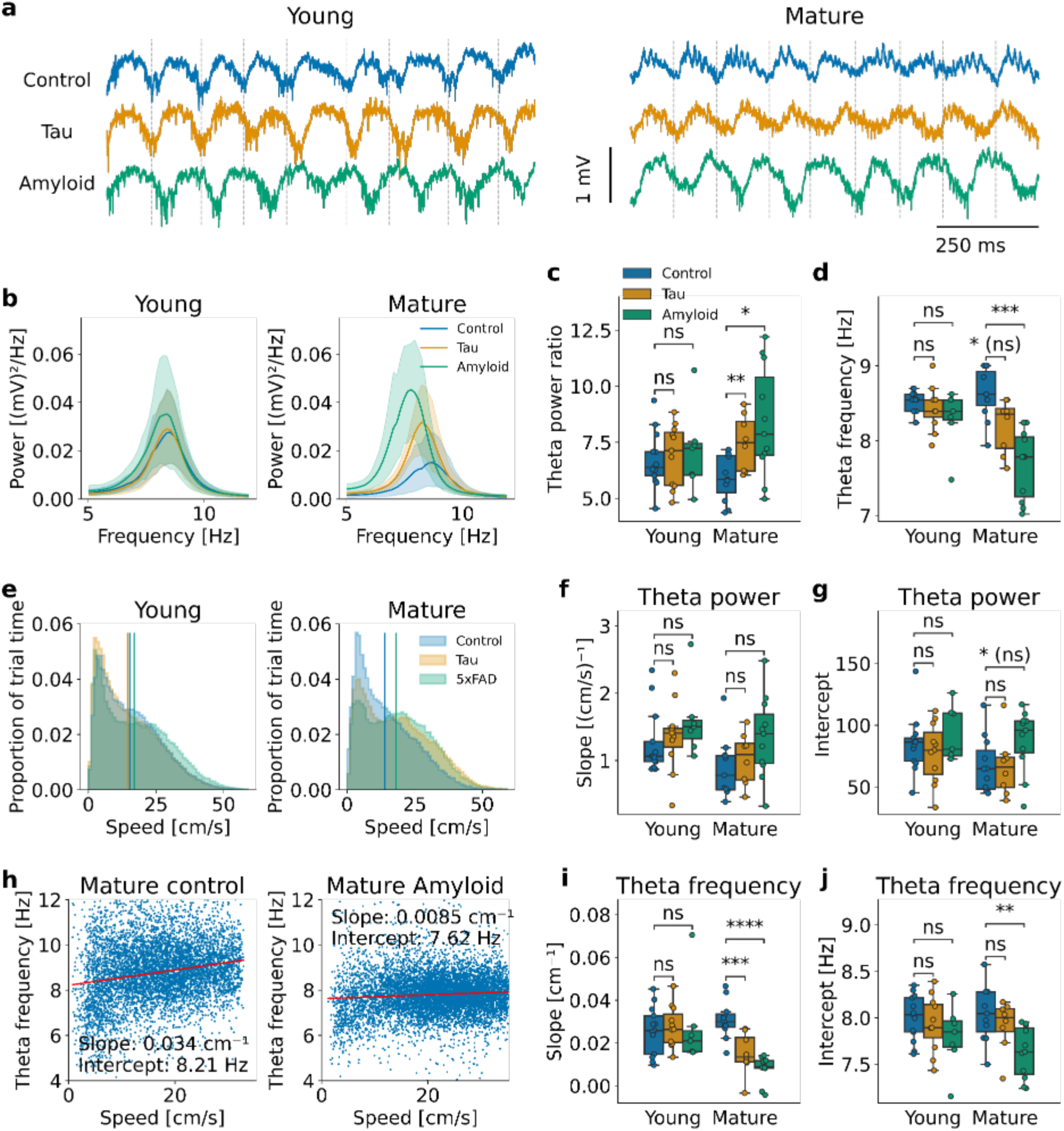
Reduced theta frequency and enhanced theta power in mature AD mice of both genotypes. (a) Examples of wide-band electrophysiological signals recorded in the MEC of control, tau and amyloid mice. Data from young and mature mice are shown separately. (b) Averaged spectrograms from all sessions. (c) Theta power ratio in the first open field trial. The theta power ratio was calculated as the power within a ±2Hz range around the theta peak compared to the combined power in the 2Hz range below and above that range. Each dot is the median of all sessions recorded in a mouse. The legend applies to (c), (d), (f), (g), (i), (j). Mann-Whitney-Wilcoxon test, two-sided with Holm-Bonferroni correction, statistical unit: mouse: young control vs. young tau: p = 0.77, U = 66; young control vs. young amyloid: p = 0.70, U = 40; mature control vs. mature tau: p = 0.0091, U = 13; mature control vs. mature amyloid: p = 0.010, U = 21 (d) Same as (c), but featuring theta frequency. Mann-Whitney-Wilcoxon test, two-sided with Holm-Bonferroni correction, statistical unit: mouse: young control vs. young tau: p = 0.27, U = 90.5; young control vs. young amyloid: p = 0.14, U = 64; mature control vs. mature tau: p = 0.020, U = 72.5 (n.s. after Holm-Bonferroni correction); mature control vs. mature amyloid: p = 0.00029, U = 116 (e) Speed distributions over trial time. Vertical lines represent the medians for each group. Mann-Whitney-Wilcoxon test, two-sided with Holm-Bonferroni correction, statistical unit: mouse: young control vs. young tau: p = 0.16, U = 96; young control vs. young amyloid: p = 0.27, U = 31; mature control vs. mature tau: p = 0.026 (n.s. after Holm-Bonferroni correction), U = 17; mature control vs. mature amyloid: p = 0.0071, U = 19 (f) Boxplots showing the slope derived from the speed-theta power ratio correlations. Each dot represents the median of all sessions recorded in a mouse. Mann-Whitney-Wilcoxon test, two-sided with Holm-Bonferroni correction, statistical unit: mouse: young control vs. young tau: p = 0.27, U = 52; young control vs. young amyloid: p = 0.067, U = 22; mature control vs. mature tau: p = 0.31, U = 31; mature control vs. mature amyloid: p = 0.066, U = 32 (g) Same as (f), but the intercept is shown. Mann-Whitney-Wilcoxon test, two-sided with Holm-Bonferroni correction, statistical unit: mouse: young control vs. young tau: p = 0.56, U = 82; young control vs. young amyloid: p = 0.64, U = 39; mature control vs. mature tau: p = 0.91, U = 46; mature control vs. mature amyloid: p = 0.049, U = 30 (h) Example scatterplots correlating the speed and theta frequency for a session of a mature control and a mature amyloid mouse. The red line represents the linear approximation; the slope and the y-intercept are printed. The speed is cropped at 90% of the max speed. (i) Boxplots showing the slope derived from the speed-theta frequency correlations. Each dot represents the median of all sessions recorded in a mouse. Mann-Whitney-Wilcoxon test, two-sided with Holm-Bonferroni correction, statistical unit: mouse: young control vs. young tau: p = 0.64, U = 63; young control vs. young amyloid: p = 0.70, U = 51; mature control vs. mature tau: p = 0.00050, U = 83; mature control vs. mature amyloid: p = 0.000082, U = 121 (j) Same as (i), but the intercept is shown. Mann-Whitney-Wilcoxon test, two-sided with Holm-Bonferroni correction, statistical unit: mouse: young control vs. young tau: p = 0.69, U = 79; young control vs. young amyloid: p = 0.21, U = 62; mature control vs. mature tau: p = 0.35, U = 56; mature control vs. mature amyloid: p = 0.0058, U = 103

It has been shown that grid cells, HD cells and border cells in MEC respond coherently to environmental manipulations ^2,3^. Thus, we hypothesized that not only grid cells, but also other spatially-tuned or directionally-tuned MEC neurons in tau mice might rotate their internal representation of space or direction in the A-B-A protocol. Indeed, already in the young cohort, HD cells and border cells in a subset of tau mice rotated their preferred firing direction between the first and the last open-field trial (**Fig. 3 a,c**, for more examples also see **Supp. Fig. 5 a**). Thus, there was a significant decrease in HD stability in mature tau mice (non-significant in young tau mice after correction for multiple testing) and in map stability for young and mature tau mice. Of note, this was also the case in young and mature amyloid mice (**Fig. 3 a,c**, for more examples also see **Supp. Fig. 5 a**). There is evidence that non-theta-tuned HD cells do not necessarily follow attractor network dynamics, but are more guided by visual cues ^51^. Thus, we wondered if the observed changes in preferred head direction were confined to non-rhythmic HD cells. For this purpose, we divided the HD cell population into theta-rhythmic and non-theta-tuned subpopulations by applying a Gaussian kernel density analysis to the distribution of theta indices of HD cells in control mice (**Supp. Fig. 6 b**, see Methods for more details). We found HD instability in both theta-tuned and non-theta-tuned HD cells (statistical significance was only reached in mature tau mice for both subpopulations, **Supp. Fig. 6 a,d,e**), and conclude that rotations in HD cells were not restricted to more visually guided non-theta-tuned HD cells.

**Fig. 6:**
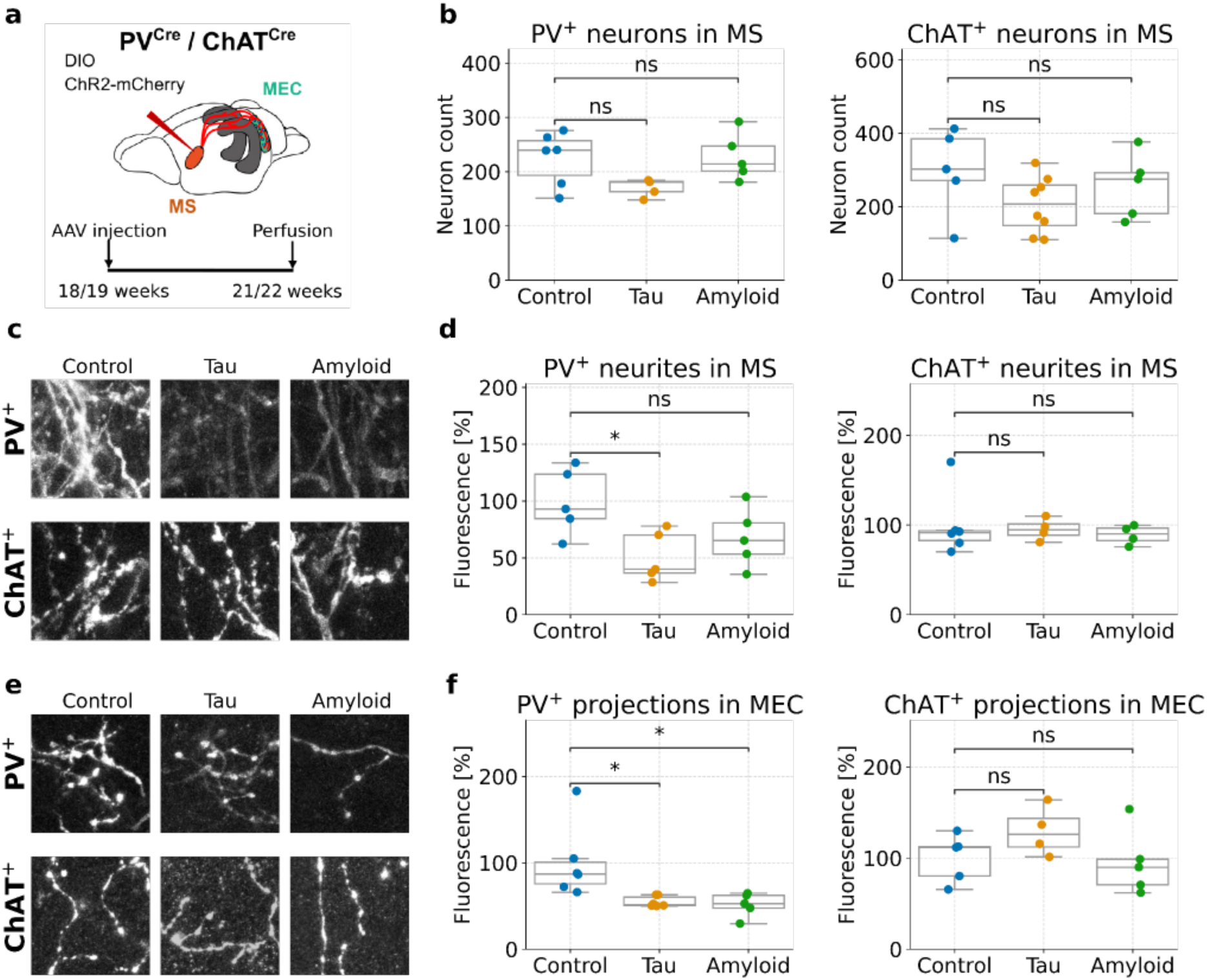
PV^+^ septal neurons exhibit a decreased density of neurites in the MS and the MEC already in young tau and amyloid mice. (a) Schematic of the experimental approach: AAV-DIO-ChR2-mCherry was injected into the medial septum (MS) of 18/19 weeks old control, tau or amyloid mice expressing Cre-recombinase in PV^+^ or ChAT^+^ neurons. Immunohistochemical analysis was performed 3 weeks later. (b) Neuron count of PV^+^ (left) and ChAT^+^ (right) neurons in the MS in control, tau and amyloid mice. Each dot indicates one mouse. One-way ANOVA, statistical unit: mouse: PV^+^: p = 0.076, F(2,13) = 3.155; ChAT^+^: p = 0.24, F(2,15) = 1.553 (c) Representative confocal images depicting mCherry^+^ neurites in the MS of PV^+^ (upper row) and ChAT^+^ (lower row) neurons in control mice, tau mice and amyloid mice. (d) mCherry fluorescent intensity in septal neurites of control, tau and amyloid mice (quantified as percentage of mCherry fluorescent intensity measured in control mice). Each dot represents one mouse. One-way ANOVA, statistical unit: mouse: PV^+^: p = 0.034, F(2,12) = 4.56, Tukey’s HSD test: control vs. tau: p = 0.029, control vs. amyloid: p = 0.17; ChAT^+^: p = 0.82, F(2,11) = 0.207 (e) Representative confocal images depicting mCherry^+^ long-range septal axons in MEC layer II of PV^+^ (upper row) and ChAT^+^ (lower row) neurons in control mice, tau mice and amyloid mice. (f) mCherry fluorescent intensity in septal long-range axons in MEC layer II/III of control, tau and amyloid mice (quantified as percentage of mCherry fluorescent intensity measured in control mice). Each dot indicates one mouse. One-way ANOVA, statistical unit: mouse: PV^+^: p = 0.014, F(2,14) = 5.877, Tukey’s HSD test: control vs. tau: p = 0.029, control vs. amyloid: p = 0.025; ChAT^+^: p = 0.24, F(2,11) = 1.606

### Rotations in grid cells, HD cells and border cells between differently-shaped environments

As we had observed unstable representations of the same spatial context across trials, we next turned to analyze the changes between the two differently-shaped environments in which local and distant cues remained unaltered (**Fig. 4 a**). Grid cells in young and mature control mice showed translation, but no rotation of their grid pattern (**Fig. 4 a-c**). In contrast, most sessions in young and mature tau mice revealed a combined phase shift and rotation of the grid cell network. Similar to the previous results, grid cells in amyloid mice did not significantly differ from those in control animals, though in particular in mature amyloid mice, the number of sessions with multiple grid cells was very low.

Furthermore, HD cells and border cells in control mice remained stable across environments, while they changed their preferred direction or shifted their firing field in a subset of sessions in young and mature tau and amyloid mice (**Fig. 4 a,d,e**, for HD cells these changes were only significant in mature tau mice).

Previous work has shown that grid cells, HD cells and border cells rotate in concert upon major changes in the environment such as the rotation of the cue card ^2,3^. It has been proposed that these cell types underlie low-dimensional attractor network dynamics. As we observed rotations across trials in these three cell types in tau mice, we wondered if these rotations were coherent. Indeed, the rotations in HD cells and border cells that were co-recorded with at least 2 grid cells proved to be consistent with the rotation of the grid cell network (**Supp. Fig. 7**). This suggests that the attractor network dynamics themselves are still intact. Since non-theta rhythmic HD cells do not necessarily follow attractor network dynamics ^51^, we compared the rotations of theta-rhythmic and non-theta-tuned HD cells that were co-recorded with “rotating” grid cells (in the case of tau mice) or border cells (in the case of tau mice and amyloid mice). While the vast majority of theta-rhythmic HD cells also exhibited rotation of their preferred firing direction, most non-theta-tuned HD cells remained stable across trials both in tau and in amyloid mice (**Supp. Fig. 8**). These results suggest that despite the intact anchoring to visual cues (as evidenced by the stable non-theta-tuned HD cells) spatial context coding in MEC is impaired in both mouse models.

**Fig. 7:**
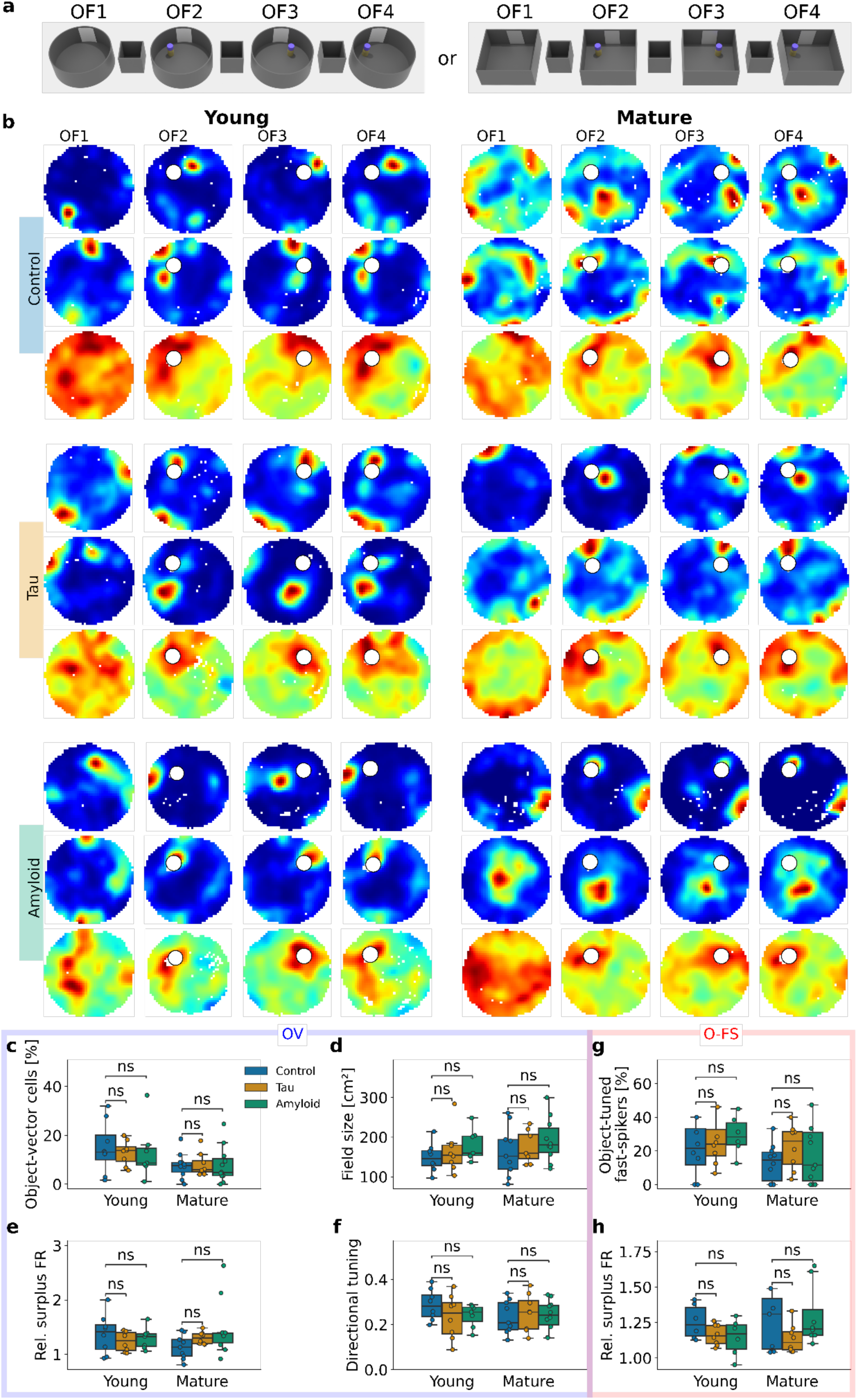
Preserved object-vector coding and object-tuned firing. (a) Schematic of the object protocol: The session started with an open-field trial (OF1) in the empty circular or square box. After a rest box trial, an object was added (OF2), which was shifted to an adjacent corner in the next open-field trial (OF3). For the last trial, the object was placed back in the original position (OF4). (b) Examples of object-vector cells (OV) and object-tuned fast-spiking neurons (O-FS) for control, tau and amyloid mice of young and mature age. For each example cell, the firing rate maps during the 4 trials of the recording protocol are shown: baseline trial, object in position 1, object in position 2, object in position 1. (c) Percentage of object-vector cells relative to the total number of recorded cells for each group. Data is grouped by mouse. The legend applies to (b)-(g). Mann-Whitney-Wilcoxon test, two-sided with Holm-Bonferroni correction, statistical unit: mouse: young control vs. young tau: p = 0.56, U = 31; young control vs. young amyloid: p = 0.35, U = 32; mature control vs. mature tau: p = 0.41, U = 43; mature control vs. mature amyloid: p = 0.42, U = 58.5 (d) Field size of object-vector cells for each group in the first object trial. In the case of several object-tuned firing fields, the largest field was taken. Data is grouped by mouse. Mann-Whitney-Wilcoxon test, two-sided with Holm-Bonferroni correction, statistical unit: mouse: young control vs. young tau: p = 0.57, U = 26; young control vs. young amyloid: p = 0.23, U = 17; mature control vs. mature tau: p = 0.66, U = 31; mature control vs. mature amyloid: p = 0.35, U = 33 (e) Ratio of the mean firing rate of object-vector cells in the area around the object in the first trial with the object compared to the baseline trial. Data is grouped by mouse. Mann-Whitney-Wilcoxon test, two-sided with Holm-Bonferroni correction, statistical unit: mouse: young control vs. young tau: p = 0.44, U = 40; young control vs. young amyloid: p = 0.78, U = 31; mature control vs. mature tau: p = 0.17, U = 21; mature control vs. mature amyloid: p = 0.13, U = 26 (f) Directional tuning of object-vector cells for each group. Directional tuning is the mean vector length of the histogram binning the firing rate of the cell around the object according to the orientation relative to the object. Data is grouped by mouse. Mann-Whitney-Wilcoxon test, two-sided with Holm-Bonferroni correction, statistical unit: mouse: young control vs. young tau: p = 0.33, U = 42; young control vs. young amyloid: p = 0.23, U = 39; mature control vs. mature tau: p = 0.81, U = 33; mature control vs. mature amyloid: p = 0.84, U = 42 (g) Percentage of object-tuned fast-spiking neurons relative to the total number of recorded fast-spiking neurons for each group. Data is grouped by mouse. For display purpose, the outliers were removed from the stripplots; the statistics were calculated including all data points. Mann-Whitney-Wilcoxon test, two-sided with Holm-Bonferroni correction, statistical unit: mouse: Mann-Whitney-Wilcoxon test, two-sided with Holm-Bonferroni correction, statistical unit: mouse: young control vs. young tau: p = 0.68, U = 28; young control vs. young amyloid: p = 0.80, U = 18; mature control vs. mature tau: p = 0.93, U = 24; mature control vs. mature amyloid: p = 0.73, U = 47 (h) Mean firing rate of object-tuned fast-spiking neurons in the area around the object in the first trial with the object compared to the baseline trial. Data is grouped by mouse. For display purpose, the outliers were removed from the stripplots; the statistics were calculated including all data points. Mann-Whitney-Wilcoxon test, two-sided with Holm-Bonferroni correction, statistical unit: mouse: young control vs. young tau: p = 0.18, U = 35; young control vs. young amyloid: p = 0.31, U = 25; mature control vs. mature tau: p = 0.46, U = 35; mature control vs. mature amyloid: p = 0.87, U = 26

**Fig. 8:**
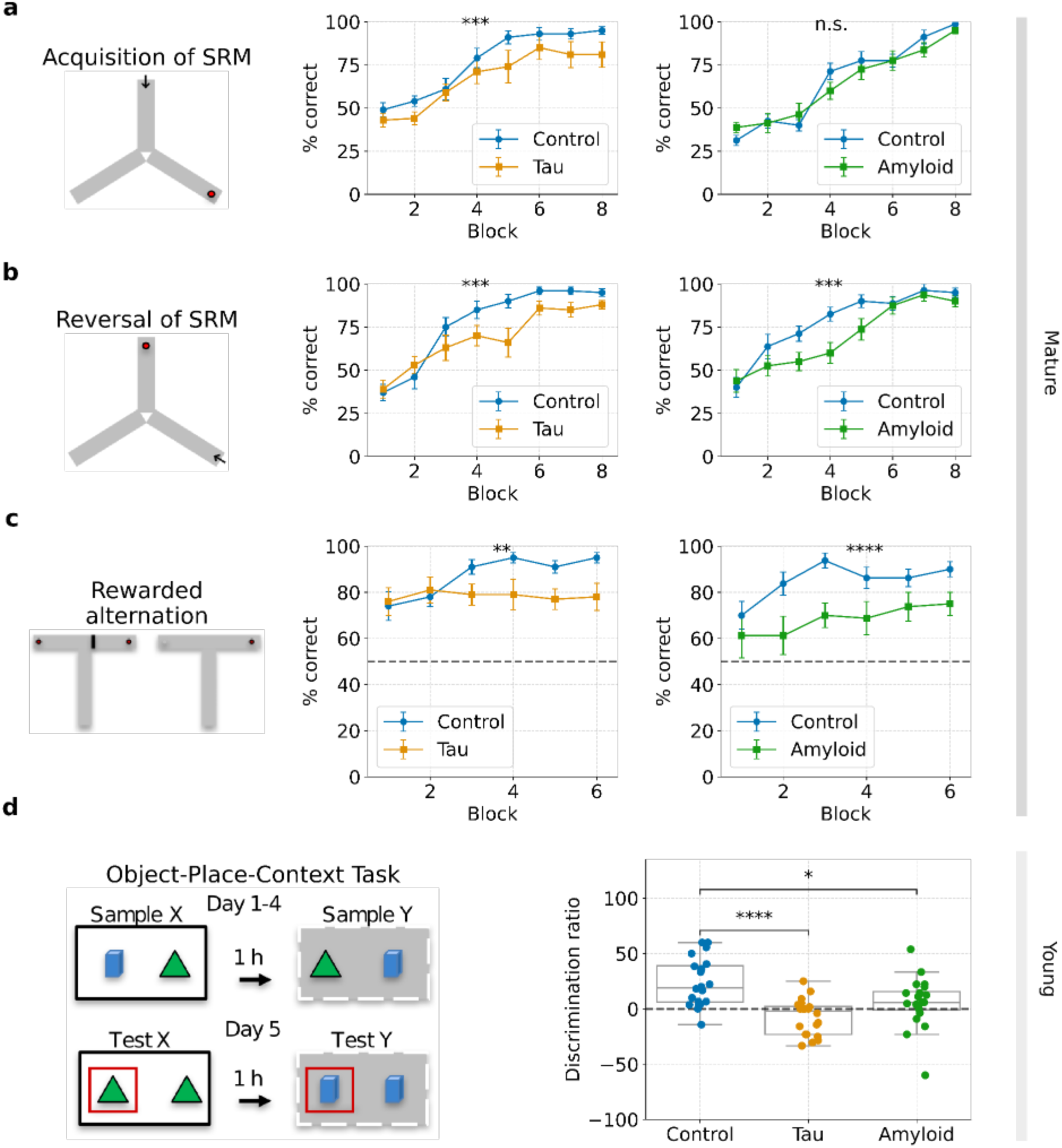
Spatial cognitive flexibility is impaired in mature AD mice, and spatial context associations are disrupted already earlier. (a) Left column: Scheme of the appetitive spatial reference memory (SRM) task on the elevated Y-maze. The red circle represents the reward and the black arrow represents the start position of the mouse. Middle column: Mean percentages of correct responses (±) SEM during the acquisition phase for mature control and tau mice (n = 10 mice per group). Each block consists of 10 trials per mouse. Two-way ANOVA: block 8: mature control 95.00 ± 2.22 %, n = 10; mature tau 81.00 ± 7.37 %, n = 10; main effect of genotype, F(1,144) = 12.14, p = 0.0007; Right column: Same for mature control and amyloid mice (n = 8 mice per group). Two-way ANOVA: block 8: mature control 98.2 ± 1.3 %, n = 8; mature amyloid 95.00 ± 1.9 %, n = 8; main effect of genotype, F(1,56) = 0.91, p = 0.34. (b) Same as (a), but the reversal learning of an appetitive spatial reference memory task on the elevated Y-maze is shown. Two-way ANOVA: block 8: mature control 95.00 ± 2.22 %, n = 10; mature tau 88.00 ± 2.49 %, n = 10, main effect of genotype; F(1,144) = 12.16, p = 0.0006; block 8: mature control 95.00 ± 2.67 %, n = 8; mature amyloid 90.00 ± 3.3 %, n = 8; main effect of genotype, F(1,56) = 12.13, p = 0.001. (c) Left column: Scheme of the non-matching to place testing on the elevated T-maze. The mouse starts in the long arm of the T-maze. In the first trial, one of the short arms containing the rewards (red circle) is blocked; in the second trial, both arms are available for the mouse. Middle column: Mean percent correct responses (±) SEM for mature control and tau mice (n = 10 mice per group). Each block consists of 10 trials per mouse. The dotted line indicates chance level performance at 50 %. Two-way ANOVA: block 6: mature control 95.00 ± 6.0%, n = 10; mature tau 78.00 ± 5.9 %, n = 10, main effect of genotype, F(1,108) = 10.77, p = 0.0014; Right column: Same for mature control and amyloid mice (n = 8 mice per group). Two-way ANOVA: block 6: mature control 90.00 ± 3.2%, n = 8; mature amyloid 75.00 ± 5.0 %, n = 8; main effect of genotype, F(1,84) = 24.24, p < 0.0001. (d) Right: Experimental protocol of the Object-Place-Context Task (OPC): The mice underwent 2 sample sessions for 4 consecutive days and two test sessions on day 5 (one in context X and one in context Y). The novel configurations in the test trials are highlighted by red frames. Left: Discrimination ratios for young control (n = 10), tau mice (n = 10) and amyloid mice (n = 10). The discrimination ratio is calculated as follows: (L2 - L1)/(L1+L2)*100, where L1 and L2 are the exploration times for object 1 and object 2, respectively. Mice naturally tend to spend more time exploring the new object-place-context configurations. One-way ANOVA: p < 0.0001, F(2,28) = 11.444. Tukey’s HSD test: young control vs. tau: p < 0.0001, young control vs. amyloid: p = 0.0236.

### Altered LFP theta oscillations in mature tau and amyloid mice

Theta oscillations provide a temporal framework for encoding spatial trajectories ^52^. When observing the electrophysiological signals of mature amyloid mice, we found a pronounced slowing accompanied by an increase in theta oscillation power (**Fig. 5 a**). Relative theta power was not only enhanced in mature amyloid mice, but also in mature tau mice, albeit to a lesser extent (**Fig. 5 b-c**). In addition, theta frequency was strongly reduced in mature amyloid mice, and the same trend could be observed in mature tau mice (**Fig. 5 d**). We wondered if these alterations in theta oscillations could be explained by different running behavior of the mice. Mature tau mice and in particular mature amyloid mice spent less time immobile, resulting in a higher overall running speed (**Fig. 5 e**). The analysis of the relationship between running speed and theta power revealed a linear relation with an unchanged slope and intercept (**Fig. 5 f-g**). This suggests that the higher theta power in mature tau and amyloid mice could be explained by the enhanced running behavior.

Conversely, when analyzing the relationship between theta frequency and running speed, we found a markedly reduced slope for mature tau and amyloid mice and for amyloid mice also a slightly diminished intercept (**Fig. 5 h-j**). Thus, theta frequency conveyed substantially less information about the current running speed than in control animals.

### Enhanced vulnerability of septal PV^+^ projections in tau and amyloid mice

Given that theta oscillatory activity is dictated by septal projection neurons targeting the MEC ^20–23^, we investigated the integrity of the PV^+^ and cholinergic (ChAT^+^) septal projections to MEC to probe for a potential mechanism of the altered LFP theta in AD mice. To this end, we injected AAV-DIO-ChR2-mCherry into the MS of 4-5 months old PV^Cre^ and ChAT^Cre^ mice (**Fig. 6 a**) and analysed the tissue 3 weeks later. The number of PV^+^ and ChAT^+^ cells in the MS was comparable between genotypes, with a tendency towards a lower neuron count in mutant mice (**Fig. 6 b**). In contrast, there was a marked reduction of the fluorescence intensity of PV^+^ neurites in the MS of tau x PV^Cre^ mice and a similar tendency in amyloid x PV^Cre^ mice, which was not the case for cholinergic neurites in ChAT^Cre^ mice (**Fig. 6 c-d**).

Furthermore, fluorescence intensity in PV^+^ projections in the MEC was reduced in mutant mice compared to the control group, whereas that of cholinergic projections remained unchanged (**Fig. 6 e-f**). Taken together, these results suggest a selective vulnerability of PV^+^ axons projecting from the MS to MEC already at early stages of AD pathology in both mouse models.

### Preserved object-vector coding and object-tuned firing

In addition to diverse spatially-tuned cells, MEC also contains object-tuned cells, specifically object-vector cells and object-tuned fast-spiking neurons. We wondered if these cell types were also impaired early in the neurodegenerative process or if the deficits were specific to spatially-tuned cells. For this purpose, on alternating days with the A-B-A protocol, the mice were recorded in the object protocol (young control: 8 mice, 74 sessions; young tau: 8 mice, 64 sessions; young amyloid: 7 mice, 52 sessions; mature control: 11 mice, 59 sessions; mature tau: 8 mice, 78 sessions; mature amyloid: 11 mice, 94 sessions) (**Fig. 7 a**). This protocol involved the introduction of an object after a standard open-field trial in the familiar environment of the mouse. Then the object was shifted to a different position and shifted back in the last trial. We found object-vector coding and object-tuned firing in all 6 cohorts (**Fig. 7 b**). The proportions of object-vector cells in tau, amyloid and control mice were similar (**Fig. 7 c**). Furthermore, basic properties such as the field size, the relative firing rate around the object compared to baseline and the directional tuning of the field at the object were comparable among groups (**Fig. 7 d-f**). Similarly, we did not observe significant differences in the proportion or relative surplus firing rate of object-tuned fast-spiking neurons (**Fig. 7 g-h**). In summary, in contrast to spatial coding, object coding was impaired neither in tau nor in amyloid mice during the observation period.

### Impaired spatial flexibility in tau and amyloid mice

To assess how the impaired spatial coding affects spatial navigation, we conducted behavioral tasks analyzing different features of MEC-dependent spatial learning and memory ^31,53,54^. First, we tested the acquisition of spatial reference memory on an elevated Y-maze in mature control, tau and amyloid mice (**Fig. 8 a**). While mature control and amyloid mice learned the task at a similar rate and reached high levels of performance, mature tau mice did not achieve the same level of performance as control animals at the end of the learning phase. To assess reversal of spatial reference memory, the goal arm on the Y-maze was subsequently switched (**Fig. 8 b**). Mature control mice quickly reached a high level of performance, whereas both tau and amyloid mice learned at a lower rate, suggesting impaired cognitive flexibility. In addition, mature tau mice exhibited a lower level of correct choices at the end of the training. Spatial working memory was analyzed using rewarded alternation (non-matching-to-place) on the T-maze ^53^ (**Fig. 8 c**). Mature control mice performed significantly better than both mature tau and amyloid mice. Of note, the performance of mature amyloid and tau mice was still clearly above the chance level of correct choices (50%) during the rewarded alternation task.

Since spatial context coding was particularly vulnerable in tau and amyloid mice already at a young age, we conducted a series of recognition memory experiments with mice of this age (5 months old). First, we employed two object exploration tasks that serve the assessment of “what” and “where” information in the context of recognition or episodic-like memory. In the sample phase of the object recognition (OR) and the object displacement (OD) task, mice encountered two identical objects in different spatial locations (**Supp Fig. 9 a+b**). After a short interval, one of the objects was replaced with an unfamiliar object at the familiar location (OR) or moved to a new location (OD) (test phase). Longer exploration times of the novel (OD) or the displaced (OR) object indicate the detection of the novel configuration. When testing young control, tau and amyloid mice in these tasks, the exploration times in the sample and the test phases did not differ (OR task: one-way ANOVA for sample phase p = 0.478, F(2,28) = 0.758, for test phase p = 0.890, F(2,28) = 0.118; OD task: one-way ANOVA for sample phase p = 0.112, F(2,28) = 2.369, for test phase p = 0.318, F(2,28) = 1.193). There was no significant impairment in the performance in mice of the three genotypes in either the OR or the OD task; however, the data suggest more variability in the performance of young tau and amyloid mice compared to the performance of young control mice during the test phase in both tasks.

Lastly, we employed an MEC-dependent object-place-context (OPC) task, which relies on the ability to form associations between the identity, the location and the spatial context of two objects (**Fig. 8 d**). Young control mice explored the novel configuration more than predicted by chance (p = 0.0004), whereas young tau and young amyloid mice explored the novel configuration at chance level (young tau p = 0.0743, young amyloid p = 0.117; Wilcoxon signed-rank test). Accordingly, there was a significant difference in discrimination ratios between the young control and the two mutant genotypes. Of note, the degree of impairment in the OPC task was similar between mature (**Supp Fig. 9 c**) and young (**Fig. 8 d**) AD mice, highlighting that altered spatial context coding in both AD mouse genotypes translates into a comparable MEC-dependent behavioral impairment already at early phases of the disease.

## Discussion

The medial entorhinal cortex (MEC) is among the first brain regions affected in Alzheimer’s disease (AD), accounting for early cognitive deficits such as spatial and episodic memory deficits ^37–42^. Various mouse models, including mice with distinct amyloid or tau pathologies, have been developed and characterized, leading to the manifestation of quite distinct impairments at the molecular, cellular and behavioral level ^43–50,55^. Our findings prompt a scenario according to which these differing pathologies ultimately converge onto a common phenotype, leading to similar effects at the cell-network and behavioral level. We found that tau and amyloid mice, starting at 5 months of age, exhibited impairments in spatial context associations, while performance on simpler spatial memory tasks, such as the object displacement test, remained intact. Corresponding to the findings at the behavioral level, HD cells and border cells in tau and amyloid mice displayed instability across recording trials, rotating their preferred firing directions or fields. Both mouse models also showed grid cell malfunction, albeit in a differential manner: While grid cells were almost absent in 8 months old amyloid mice, the grid cell network in tau mice became unstable, showing translation and rotation of the grid map across trials. Furthermore, we found that in contrast to spatial coding, object-coding in MEC remained intact throughout the observation period. Additional commonalities included reduced theta frequency, elevated theta power and disrupted theta speed coding. At the cellular scale, we identified selective degeneration of septal parvalbumin (PV^+^) projections to the MEC as a common feature, potentially underlying the here-identified impairments at the cell network level. Together, these results highlight how different pathologies in AD-related neurodegeneration converge onto similar manifestations of disrupted spatial and temporal coding within the MEC, culminating in impaired spatial context coding as a shared behavioral phenotype.

Previous studies that addressed grid cell coding in mouse models of amyloid or tau pathology reported severe disruptions of grid cells at moderate to advanced stages of neurodegeneration ^47–50^. In the present study, we demonstrate that in Tau P301S mice, the grid cell malfunction primarily manifested as an inability to reflect the spatial context, while the grid cell patterns themselves remained largely preserved during the observation period. In contrast, in mature 5xFAD mice, we observed a pronounced grid cell reduction in line with previous findings. As we did not detect differences in the spatial stability of the grid cell network between young control mice and young amyloid mice, it is likely that instability in spatial coding of border and head-direction (HD) cells emerges concurrently with the onset of severe grid cell loss in this mouse model. In line with this hypothesis, grid scores in non-fast-spiking cells were reduced already in young amyloid mice. Moreover, the remaining grid cells in mature amyloid mice displayed enlarged firing fields and increased grid spacing, indicative of reduced spatial precision. Together, these findings suggest that grid cell dysfunction represents a convergent feature of MEC impairment in the context of neurodegeneration.

The earliest shared impairment observed in both mouse models was a disruption in spatial context coding. Beginning at 5 months of age, HD and border cells in both tau and amyloid mice failed to maintain stable preferred firing directions or fields across recording trials, despite unaltered local and distal cues. Previous work has shown that in 7–13 months old amyloid model mice, MEC neurons were impaired in remapping between two different linear tracks ^47^. In contrast, our findings indicate that a stable representation of the same environment could not be maintained. Interestingly, a subset of non-theta-tuned HD cells in both tau and amyloid mice remained stable across trials, despite co-recorded cells displaying instability. This observation suggests that the loss of stability is not due to degraded sensory input, as anchoring to visual cues in the case of visually driven HD cells appears to remain intact. Furthermore, we observed coherent rotations of grid cells, (theta-tuned) HD cells, and border cells, indicating that the low-dimensional attractor dynamics that coordinate these spatial cell types remain intact. This supports the hypothesis that the network architecture underlying these interactions is hard-wired and thus resilient to the incipient pathologies.

The observed instability in spatially tuned neurons likely underlies the behavioral deficits in spatial context processing seen in both tau and amyloid mice. While performance in simple spatial memory tasks, such as the object displacement test, remained intact, both models exhibited impairments in more complex tasks requiring the integration of spatial cues across time and contexts, such as the object-place-context task. This behavioral dissociation mirrors the neural data: Although individual spatial cell types retained basic firing properties, their instability across trials likely reflects the lack of formation of coherent spatial representations necessary for contextual memory. The coherent rotation observed across grid, HD and border cells suggests that the attractor network dynamics remain intact but fail to reliably anchor to external spatial cues. This partial preservation of network coordination, combined with impaired cue anchoring, may explain why spatial computations dependent on internal dynamics remain functional, while tasks requiring stable, context-specific representations are impaired.

In contrast to the instability observed in spatially modulated cells, object-vector (OV) cells in both tau and amyloid mice remained intact throughout the observation period. We also identified stable object-tuned fast-spiking neurons, suggesting that object-related firing in the MEC is generally preserved. Notably, OV cell function in the MEC in the context of Alzheimer’s disease mouse models has not been studied so far. Our results suggest that object-related representations are more resilient to early neurodegenerative processes than spatial context coding. It is tempting to speculate that the resilience of OV cells at incipient stages of AD may well account for the finding that landmark-based orientation is better preserved than non-landmark-based navigation in patients with early AD ^56,57^. This identifies object-vector coding as a putative target for alternative strategies to be leveraged to improve navigation in patients with early Alzheimer’s disease. For instance, Davis and colleagues found that augmenting salient cue information enhanced performance in AD patients ^56^.

Both tau and amyloid mouse models showed altered LFP theta oscillations, characterized by lower theta frequency and increased theta power. These changes are particularly relevant given the central role of theta rhythm in coordinating temporal coding across hippocampal-entorhinal circuits. Interestingly, similar changes in theta power have been observed in humans with mild cognitive impairment (MCI) ^58,59^ and dementia ^60,61^. The higher running speed in mature amyloid mice, and the same tendency in mature tau mice, is most likely reflected in the elevated theta power in both genotypes. This is in line with the notion that theta amplitude is controlled by cholinergic neurons in the medial septum ^22^ and with our finding that the integrity of septal cholinergic projections to the MEC is preserved.

Alternatively, higher theta power could represent a compensatory mechanism masking underlying circuit instability. In contrast, theta frequency is controlled by the GABAergic interneurons in the medial septum ^62,63^. Together, these findings highlight how alterations in theta dynamics can disrupt the temporal structure of MEC activity, potentially contributing to deficits in both navigation and episodic memory functions.

In the context of altered theta oscillations in both AD mouse models, it is worth emphasizing that most AD studies in mouse models, but also in human, homed in on different subregions of the hippocampal formation, i.e. the entorhinal cortices and hippocampus proper. It stands to reason that these regions obtained the highest attention as pathology commences here ^37^. We extended our investigations to the MS as it is considered the pacemaker of the entire hippocampal formation. Indeed, we and others found that by virtue of the specific connectivity of septal PV^+^ projection neurons to GABAergic interneurons in the target regions ^21,26^, these neurons are ideally suited to orchestrate network activity in the entire hippocampal formation. Thus, septal PV^+^ cell-generated theta oscillations are at the core of the local LFP that constitute a prerequisite for the ordered firing of the different cell types within a theta cycle ^64^. It is tempting to speculate that the more pronounced disruption of theta-rhythmic HD cells in comparison to non-theta-tuned HD cells reflects this upstream pathology, i.e. the vulnerability of theta-generating septal PV^+^ cells. Furthermore, the instability of MEC spatial representations is very likely related at least in part to malfunction of the septal pacemaker. Together these considerations point to a pathology upstream of the MEC/hippocampal formation that is worth considering for further therapeutic interventions.

Notably, our immunohistochemical results revealed a degeneration of septal PV^+^ projections to the MEC, but not cholinergic projections, at the stages analyzed here. This is interesting in the context of human studies that emphasized the vulnerability of cholinergic neurons in the basal forebrain at incipient phases of the disease ^65, 66, 67,68^. The enhanced vulnerability of cholinergic projections has been attributed hypothetically to their long axons and complex branching that likely denote a high metabolic need ^66^. Thus, our results strongly indicate that, at least in mouse models of AD, axons of GABAergic fast-spiking cells such as long-ranging PV^+^ septal projections exhibit enhanced vulnerability in neurodegenerative processes before cholinergic projections are affected. Indeed, electrophysiological recordings from septal PV^+^ neurons in freely moving mice allow the inference that this cell population must have a high metabolic rate, given that 40% exhibited fast spiking (unpublished data).

In summary, our study reveals that despite differing underlying pathologies, both tau and amyloid mouse models converge on shared disruptions in spatial and temporal coding within the MEC, resulting in a common behavioral phenotype characterized by impaired spatial context processing. These deficits emerge early in the course of the disease, highlighting the vulnerability of MEC circuits at the incipient stages of neurodegeneration. By identifying convergent network-level dysfunctions across models, our findings raise the possibility that early alterations in spatial coding could serve as a sensitive functional biomarker for AD, potentially paving the way for the development of simple diagnostic tools targeting MEC-related navigation and memory functions.

## Methods

### Subjects

For this study, electrophysiological recordings were performed in a total of 68 mice, 61 of which were included in the analysis. 5 mice were excluded because the position of all shanks could either not be retrieved (1 mouse) or found to be outside the superficial layers of MEC (4 mice). 2 mice were excluded because the results of the A-staining were inconsistent with the documented genotype/age. The remaining 61 mice consisted of 13 4-7 months old and 11 7-11 months old male wild-type mice maintained on a C57BL/6N background, 11 4-7 months old and 8 7-10 months old male Tau P301S mice and 7 4-7 months old and 11 7-11 months old male 5xFAD mice. Mice were divided into “young” and “mature” cohorts based on their age in the first recording session (4-6 months and 7-9 months, respectively).

For the behavioral experiments, a total of 67 mice were used, trained in 3 separate groups: The first group consisted of 10 6-8 months old male wild-type mice and 10 6-8 months old Tau P301S mice. These mice were tested on the acquisition of spatial reference memory on the Y-maze, the reversal learning on the Y-maze, followed by rewarded alternation on the T-maze and the object-place-context task. The same order of experiments was performed by an age-matched group consisting of 8 6-8 months old male wild-type mice and 8 6-8 months old 5xFAD mice. Both groups are referred to as “mature” cohorts. The third group (“young” cohorts), consisting of 11 5 months old wild-type mice, 10 5 months old Tau P301S mice and 10 5 months old 5xFAD mice, performed the object recognition task, the object displacement task and the object-place-context task before the mice reached 6 months of age.

All mice were kept on a 12 h light/dark cycle. All experiments were performed during the light period. Mice that were implanted with a silicon probe were single-housed. Food was restricted to maintain a body weight of 85% of the initial body weight with food ad libitum, whereas mice had water ad libitum. All procedures followed the European Council Directive (86/609/EEC) and were approved by the Governmental Supervisory Panel on Animal Experiments of Baden Württemberg in Karlsruhe (G-234/19).

### Surgical procedure

For the electrophysiological recordings, the mice were implanted with either one 64-channel H64LP NeuroNexus probe or one 64-channel H10 Cambridge Neurotech probe. One old control mouse was implanted bilaterally with 64-channel H10 Cambridge Neurotech probes. Before implantation, the probes were mounted on custom-made microdrives, which allowed for linear movement in the dorsoventral axis. The mice were anesthetized with 1.5-3% isoflurane before they were fixed to a stereotactic frame. During the surgery, anesthesia was maintained using 0.5-1.5% isoflurane. After exposure of the skull, 2 miniature screws were attached to the skull. While one screw served as a fixation screw, the other screw, which was inserted above the cerebellum, was used to ground the electrophysiological signal. A craniotomy was performed above the implantation site and the transverse sinus was exposed. The probe was implanted 3.1 mm from the midline, 0.2 mm anterior of the transverse sinus at an angle of 6-7° in the sagittal plane, with the probe tip pointing slightly in the posterior direction. The microdrive was fixed to the skull with dental cement at a depth of 0.5-0.7 mm. After surgery and during the next 72 h, the mice were injected with carprofen (0.1 mg/kg BW) s.c for analgesia and given at least 6 days of recovery before food restriction and behavioral training.

### Recording setup

For the electrophysiological recordings, a square environment (70 cm x 70 cm, height of walls approx. 30 cm) and a circular environment (diameter of 80 cm, height of walls approx. 35 cm) were used. Both arenas were black and had a white cue card attached to one wall. Assuming the cue card pointed north, the mouse would perceive metal bars and black curtains in the north, east and south direction. The opening of the recording setup would point west, giving view to the cabinet where the mouse cages were kept. A pellet dispenser (CT-ENV-203–5 pellet dispenser, Med Associates) was mounted above the center of the arena and controlled by a microcontroller (Arduino Uno). Additionally, there was a camera attached to the metal bars above the arena. The light in the room was dimmed.

### Behavioral training and recording protocols

One day after the initiation of food restriction, the mice were alternately assigned to either the square environment or the circular environment, called the “familiar environment” hereafter. The assignment was done per group. The behavioral training was performed only in the assigned arena. During the training, the mouse randomly foraged for food (AIN-76A Rodent Tablet 5 mg; TestDiet), which was automatically dispensed at a rate of 1-3 pellets/min, from the pellet dispenser. Training sessions lasted for 20-30 min and were performed 1-2 times per day. On the second or third training day, the mouse was connected to the recording system. The probe was gradually lowered (in steps of ∼62.5 µm for NeuroNexus probes or ∼125 µm for Cambridge Neurotech probes). Once a complete coverage of the environment was reached within 20 min and the electrophysiological recordings showed theta activity and spikes on at least one shank, the mouse was transferred to the recording phase.

Mice performed two different protocols on alternating days: the A-B-A protocol and the object protocol. The A-B-A protocol consisted of 3 open-field trials interspersed with 2 rest trials. The first open-field trial was conducted in the familiar environment and involved random foraging for 20 min. After that, the mouse was put in a rest box (30 cm x 30 cm) for 10 min. The rest box was placed at the center of the recording environment. Next, the box was replaced by a differently shaped environment, i.e. the circular arena if the familiar environment was the square arena and vice versa. The cue card in the novel environment was attached to the wall in the same direction as in the familiar environment. Thus, all local and distant cues were kept unaltered. Another random-foraging trial was conducted for 20 min. After another rest box trial, a third open-field trial in the familiar environment was conducted for another 20 min. Again, all local and distant cues remained unchanged. After each training or recording session, all boxes were cleaned with ethanol.

The object protocol consisted of 4 open-field trials, all conducted in the familiar environment. During the first open-field trial, the mouse randomly foraged for food in the familiar arena for 20 min. After a 10 min rest trial, an object was introduced in the arena and the mouse performed another random-foraging trial for 20 min. The object consisted of a 100 ml bottle (diameter 7 cm) with a dark blue lid, filled with sand. Four transparent 2 ml tubes were glued on the lid to prevent the mice from climbing on the object. The object was placed in the northwest corner (20 cm north and 20 cm west from the middle of the arena). After another 10 min rest trial, the object was shifted to the northeast corner (20 cm north and 20 cm east from the middle of the arena) for the third random-foraging trial. Finally, after another 10 min rest trial, a fourth random-foraging trial with the object shifted back to the northwest corner was performed.

When the mouse had completed both protocols, the probes were lowered by ∼62.5 µm (NeuroNexus probes) or ∼125 µm (Cambridge Neurotech probes). In the case of the bilateral implant, only one probe was lowered, alternating between the right and the left side.

### Electrophysiological recordings

The mouse was connected to a data acquisition system (RHD2000-Series Amplifier Evaluation System, Intan Technologies, analog bandwidth 0.09–7603.77 Hz, sampling rate 20 kHz) by a lightweight cable. We used custom-written software (https://github.com/kevin-allen/ktan) to control the recording process. The data were then processed in a semi-automated manner using Kilosort (https://github.com/jamesjun/Kilosort2) and Phy (https://github.com/cortex-lab/phy). To ensure cluster quality, only clusters with a distinct waveform, > 100 spikes in the spike time autocorrelation (from 0 to 30 ms), a mean firing rate > 0.1 Hz and a refractory period ratio < 0.15 were included in the analysis. The refractory period was defined as the quotient of the mean number of spikes in the bins 0 to 1.5 ms and the maximal number of spikes in any bin between 5 and 15 ms (using a spike time autocorrelation from 0 to 25 ms and a bin size of 0.5 ms).

The position of the mouse was tracked using 3 differently-colored LEDs attached to the head of the animal. The light signal was recorded by a video camera above the arena at a frame rate of 25 Hz.

### Histology

When the electrophysiological signal indicated that all shank tips had reached layer I or when the probe could not be lowered further into the brain, the experiment ended, and the mouse was deeply anesthetized by an intraperitoneal injection of ketamine (20%; 50 mg/ml) and xylazine (8%; 20 mg/ml). The mouse was transcardially perfused using phosphate-buffered saline (PBS) and 4% paraformaldehyde (PFA). After extraction of the brain, the brain was stored in PFA for 24 h. Then, the brain was washed with PBS, and the posterior part of the right hemisphere was cut into sagittal slices (thickness of 50 µm) using a vibratome (Leica Germany). The brain slices were stained with Cresyl violet and mounted on slides, which were digitized with a motorized widefield slide scanner (Axio Scan.Z1, Zeiss).

### AAV injections

AAV-double floxed-hChR2(H134R)-mCherry-WPRE-pA (AAV DIO ChR2-mcherry) virus was a gift from Karl Deisseroth and produced from Addgene (USA). This virus carries an inverted version of Channelrhodopsin 2 fused to the fluorescent marker mCherry. In the presence of Cre recombinase, the cassette is inverted into the sense direction and the fused proteins are expressed from the EF1 promoter.

We injected AAV DIO Chr2-mCherry into the MS of 18 to 19 weeks old male ChAT^Cre^ (ChAT tm2(cre)Lowl/J, purchased from the Jackson Laboratory) (6 mice), Tau P301S x ChATCre (8 mice), 5xFAD x ChAT^Cre^ (5 mice) or PV^Cre^ ^69^ (6 mice), Tau P301S x PV^Cre^ (6 mice), 5xFAD x PV^Cre^ mice (5 mice).

Animals were anesthetized with isoflurane, mounted in a stereotactic apparatus and kept under isoflurane during surgery. For MS injections a small craniotomy was made 1 mm anterior to bregma. The AAV DIO Chr2-mCherry was delivered by a glass micropipette with a tip resistance of 2 to 4 MΩ, 4 mm below the cortical surface into the MS. 200 nl of recombinant AAV DIO Chr2-mCherry were injected. The scalp incision was sutured, and post-surgery analgesics were given to aid recovery (0.03 mg/kg Metamizol).

Immunohistochemical experiments were done 3 weeks after injection. We did not analyse mice in which the injection site did not correspond to the targeted area.

### Immunohistochemistry

Mice were transcardially perfused wit 4 % paraformaldehyde. Coronal sections (from MS) and sagittal sections (from MEC) were cut at 50 µm thicknesses on a vibratome and washed with phosphate buffered saline (PBS). For analysing fluorescent intensity of mCherry in the MS, the sections were mounted on 0.1 % gelatin-coated glass slides and mounted in Mowiol 40-88.

For visualizing long-range projections in MEC free-floating sections were permeabilized and blocked for 2 hrs with PBS containing 5% bovine serum albumin (BSA) and 0.2 % Triton X-100. The incubation of the sections with dsRED antibody (rabbit, 1:1000) was performed for 24 h at 4°C. Sections were washed with PBS and incubated for 2 h with Cy3-conjugated secondary antibody (1:1000 Jackson Immunoresearch, Newmarket, UK). After repeated washing with PBS sections were mounted on 0.1 % gelatin-coated glass slides and mounted in Mowiol 40-88.

Confocal images were taken with a 63x objective using a Zeiss LSM 700 microscope (Zeiss, Jena, Germany). Three sections were analysed from each mouse (at least 150 µm apart from each other). Images of eight 30 x 30 µm ROIs (regions of interest) were taken from each section and fluorescent intensity analysed using the Fiji software.

For counting the neuron numbers in control and mutant mice, 6 wild-type mice, 9 Tau P301S mice and 7 5xFAD mice of 5 months of age and 8 wild-type mice, 7 Tau P301S mice and 9 5xFAD mice of 8 months of age were transcardially perfused as described above. Sagittal sections were cut at 50 µm thickness and washed with PBS. The sections were blocked with BSA (5%) and Triton X-100 (0.2%) for 2 h. Then, they were incubated with a primary antibody (monoclonal anti-NeuN (1:2000, Millipore, Temecula, CA, USA) for 24 h at 4°C. After washing with PBS, the sections were incubated with a secondary antibody (Cy3, Jackson Immunoresearch, 21 Newmarket, UK, 1:1000). After repeated washing with PBS, the sections were mounted on 0.1% gelatin-coated glass slides and mounted in Mowiol 40-88. The sections were analyzed by confocal microscopy (Zeiss LSM 700). For each animal, the stained neurons in layer II and layer III were counted manually in 5 sections spanning the superficial layers of MEC (each 150 µm apart). The results were averaged per mouse.

### Behavioral experiments

#### Acquisition of spatial reference memory on the elevated Y-maze

Spatial reference memory on the elevated Y-maze was examined as previously described ^70^. The Y-maze was made of black painted wood and had a central polygonal area (diameter of 14 cm) to which three arms were attached (50 x 9 cm, surrounded by a 0.5 cm high beading). A plastic food well was located 5 cm from the distal end of each arm. The maze was located in a room unfamiliar to the mice, with prominent distal cues. It was elevated 80 cm above the ground on a central stand on which the entire maze could be rotated to prevent the use of intra-maze cues. The mice were familiarized with the maze until they were running freely on the maze and readily consuming sugar pellets from the food wells. A target arm (defined according to its given spatial location relative to the room cues) was designated for each mouse; here, the mouse would receive a reward of 3 sugar pellets. Target arms were counterbalanced with respect to genotype such that approximately equal numbers of mutant and control animals were trained to each of the arms. The start arm for each trial was determined by a pseudorandom sequence with equal numbers of starts from each arm in any session and no more than three consecutive starts from the same arm. The testing period lasted for 8 consecutive days, each mouse performing 10 trials per day.

#### Reversal of spatial reference memory on the elevated Y-maze

One day after finishing the acquisition of spatial reference memory, a new target arm (defined according to its given spatial location relative to the room cues) was designated for each mouse. Target arms were again counterbalanced with respect to genotype such that approximately equal numbers of mutant and control animals were trained to each of the arms. The start arm for each trial was determined by a pseudorandom sequence with equal numbers of starts from each arm in any session and no more than three consecutive starts from the same arm.

#### Spatial working memory on the elevated T-maze

Spatial working memory was assessed on an elevated T-maze ^70^. First, the mice were habituated to the maze and to receiving sugar pellets over several days before spatial non-matching-to-place testing. Each trial consisted of a sample run and a choice run. On the sample run, the mice were forced either left or right by the presence of a wooden block, according to a pseudorandom sequence (with equal numbers of left and right turns per session, and not more than two consecutive turns in the same direction). A reward of 3 sugar pellets was available in the food well at the end of the arm. The block was then removed, and the mouse was placed at the end of the start arm, facing the experimenter. The delay interval between the sample run and the choice run was about 10-15 sec. The animal was rewarded for choosing the previously unvisited arm (i.e. for alternating). Entry into an arm was defined when a mouse placed all four paws into that arm. Mice were run one trial at a time with an inter-trial interval of about 10 min. Mice performed 4 trials per day over 10 days of testing.

#### Object-Place-Context (OPC) Task

The OPC task was performed by the same mice that underwent the spatial working memory task on the T-maze (8 months old mice) or 4 days after the OR task (5 months old mice)..

To learn the context-place-object associations, the mice were first exposed to a pair of distinct objects (A1 and B1) in context X, and an identically looking pair of objects (A2 and B2) whose position was reversed in context Y. Objects A and B were of similar size, but differed in colour, shape and surface texture. The arenas had a size of 50 x 30 x 18 cm each; one arena was equipped with black walls and a white floor (context X), the other arena with grey-and-white striped walls and a grey floor (context Y).

The mice were exposed to each context for 10 min per day for the first three days, and for 5 min on day 4. The order of exposure to the two contexts alternated daily (e.g. X-Y, Y-X, X-Y, Y-X), and on each day the two exposures were separated by approximately 1 hour.

On the 5th day, the mice performed a 5 min test trial in each context. In the test trials, one of the objects was replaced by a copy of the other object (e.g. A1 by B2 in context X and B2 by A1 in context Y), so that the mice were exposed to two identical objects (e.g. B2 and B1 in context X or A1 and A2 in context Y). Each test trial was separated by approximately 1 hour. The identity of objects A and B, the location of the object that had not previously been paired with the test context and the order of trials in context X and Y were counterbalanced for each genotype. The object, which was unfamiliar in a location for a given context, was defined as “novel”. A discrimination index was calculated as 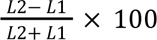, where L1 and L2 are the exploration times for the “familiar” object and the “novel” object, respectively.

#### Object Displacement (OD) Task

For the OD task ^71^, mice were individually habituated to an empty grey square arena (45 x 45, height of 25 cm) for 3 days. For the sample phase, two identical copies of an object were placed in the arena at a distance of 15 cm from each other, and mice were allowed to explore the objects for 10 min. After an inter-trial interval of 5 min, mice were returned to the arena for a test phase of 5 min. During the test phase, one object was at a new position that was at a distance of 20 cm from the previous location. On each trial in the sample and the test phase, mice were placed into the arena at the same starting point, facing away from the objects. The arena and objects were wiped with 70 % ethanol between trials to minimize olfactory cues. Exploration was defined as sniffing or touching the object with the nose or forepaws. A discrimination index was calculated as 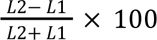, where L1 and L2 are the exploration times for the non-displaced object and displaced object, respectively.

#### Object Recognition (OR) Task

Two days after completion of the OD task, an OR task was performed ^72^. Mice were individually habituated to an empty grey rectangular arena (50 x 30, height of 18 cm) for 3 days. For the sample phase, two identical copies of an object (A1 and A2) were placed at a distance of 15 cm from each other, and mice were allowed to explore the objects for 10 min. After an inter-trial interval of 5 min, the mice were returned to the arena for a test phase of 5 min. During the test phase, the arena contained a third identical copy of the objects used in the sample phase (A3) and a novel object (B1). The objects (A3 and B1) were placed in the locations previously occupied by the objects in the sample phase (A1 and A2). On each trial in the sample and the test phase, the mice were placed into the arena at the same starting point, facing away from the objects. Exploration was defined as sniffing or touching the object with the nose or forepaws (like in the OD task). A discrimination index was calculated as 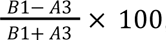, where B1 and A3 are the exploration times for the new object and the familiar object, respectively.

### Analysis of the electrophysiological recordings

#### Firing rate maps

Firing rate maps were constructed by dividing the recording environment into bins of 2 cm x 2 cm. The occupancy map calculated from the time spent in each bin was smoothed with a Gaussian kernel with a standard deviation of 2 cm. The firing rate map was then calculated by dividing the number of spikes in each bin by the time spent there. For better visualization, the firing rate maps shown as examples were calculated using a Gaussian kernel with a standard deviation of 4 cm.

#### Grid score

The grid score was calculated from the spatial autocorrelation of the firing rate map. If 6 fields around the center of the spatial autocorrelation map were detected, a doughnut-shaped region containing these fields was defined. This region was then rotated by multiples of 30° and correlated with the non-rotated region. The grid score was defined as 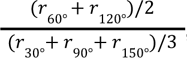, where  is the Pearson correlation coefficient between the non-rotated and rotated regions.

#### Information score

For the calculation of the information score, the firing rate map was re-calculated without smoothing. The information score was defined as 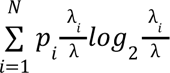, where  is the number of bins of the firing rate map,  is the probability of the occupancy bin , λ is the mean firing rate and λ is the firing rate in the bin . This definition follows the work of Skaggs et al. ^11^.

#### Border score

The border score calculation was adapted from the paper by Solstad et al. ^3^, using the following formula: 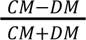. is defined as the maximal proportion of field pixels that were border pixels belonging to one wall across all field x wall combinations.  is the normalized mean closest distance of the field pixels to the closest wall weighted by the firing rate of the pixel.

First, all firing fields were detected in the firing rate map. In this context, a firing field was defined as a conglomeration of at least 20 contiguous pixels with a minimal firing rate of 4 Hz in at least one of the pixels and at least 30% of the peak firing rate of the whole firing rate map. In the next step, all border pixels were detected. For the circular environment, 36 overlapping wall segments spanning 45° each were defined.

#### Head-direction score (HD score) and directional tuning distributive ratio

To calculate the HD score, a firing rate histogram per head direction was obtained using a bin size of 10° and a Gaussian kernel with a sigma of 10° for smoothing. The HD score was defined as the mean vector length of this histogram.

The directional tuning ratio was calculated as described in the work of Kornienko et al. ^51^ in order to distinguish between pseudo-HD selectivity due to spatial selectivity and true directional selectivity. In brief, first, the head-direction occupancy histogram was obtained for each pixel of the firing rate map. Then the predicted HD selectivity solely based on the spatial selectivity *R_Pred_* was calculated as follows: 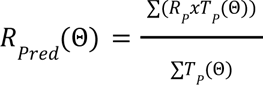where *R_P_* is the firing rate in each pixel of the spatial firing rate map and *T_P_* (Θ) is the time spent looking in the direction Θ whilst in this pixel. Finally, the distributive ratio *DR* was defined as the similarity between the observed (*R_obs_*) and predicted HD tuning curve: 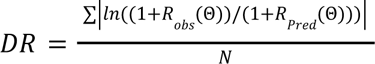 where *N* normalizes the term by the number of bins in the HD tuning curve.

#### Cell type classification

MEC neurons were classified hierarchically, i.e. one cell could also be assigned to one cell type in the following order: Fast-spiking neurons > grid cells > border cells > HD cells > others. Cells had to fulfill the respective criteria for the first **and** the last open-field trial in the A-B-A protocol to ensure stability across the entire recording session. Shuffling thresholds were generated by repeating the calculation 200 times after shifting the positional data (minimal shift of 20 sec).

Fast-spiking neurons were defined as neurons with a mean firing rate > 10 Hz.

Grid cells were defined as neurons that did not qualify as fast-spiking neurons and had a grid score and an information score exceeding the 95th percentile of the respective random distribution in the first and the last open-field trial in the A-B-A protocol.

Border cells were defined as neurons that did not pass the criteria for fast-spiking neurons or grid cells, but had a border score exceeding the 95th percentile of the shuffled distribution (but at least 0) and an information score exceeding the 95th percentile of the shuffled distribution (but at least 0.5) in the first and the last open-field trial in the A-B-A protocol.

HD cells were defined as neurons that did not pass the criteria for fast-spiking neurons, grid cells or border cells, but had an HD score exceeding the 95th percentile of the shuffled distribution, a peak directional firing rate of at least 4 Hz and a directional tuning distributive ratio > 0.2 in the first and last open-field trial of the A-B-A protocol.

Object-vector cells and object-tuned fast-spiking neurons were defined based on their firing behavior in the object protocol: Object-vector cells were defined as MEC neurons with a mean firing rate < 10 Hz in the first open-field trial, a grid score not exceeding the 95th percentile in the first open-field trial, at least one new firing field around the object in the second open-field trial with a distance > 4 cm from the object center, a Pearson r correlation of the area around the object in the second and third open-field trial > 0.4, an information score exceeding the 95th percentile in the second and third open-field trial and a peak firing rate around the object location > 2 Hz in the second and third open-field trial. This definition is adapted from the work of Høydal et al. ^27^. The area around the object was defined as a circle with the object at its center and a radius of 20 cm.

Object-tuned fast-spiking neurons were defined as neurons with a mean firing rate > 10 Hz in the first open-field trial, an information score exceeding the 95th percentile in the second and third open-field trial and a mean firing rate around the object exceeding the mean firing rate in the trial by at least 10% in the second and third open-field trial. The concept of object-tuned fast-spiking neurons is derived from the paper by Caputi et al. ^28^.

#### Grid spacing and grid module separation

The grid spacing was calculated from the spatial autocorrelation of the firing rate map of grid cells. First, the peaks in the spatial autocorrelation were detected. After removal of the peak at the center, the grid spacing was obtained by averaging the distances of the remaining 6 peaks from the center.

To separate different grid modules, for sessions with at least 2 co-recorded grid cells, we performed a kernel density estimation of the distribution of grid spacings, using a Gaussian kernel with a bandwidth of 3 cm. Next, we obtained the relative extrema of the fitted function. In case a trough was found, the session contained 2 grid modules and the trough determined the cutoff between them. We did not record sessions with more than 2 modules.

#### Grid cell rotation and phase shift

Calculation of grid cell rotation and phase shift was attempted for sessions with at least 2 co-recorded grid cells. For each grid cell, the z-scored firing rate map was calculated (bin size = 2 cm, smoothing kernel = 2 cm) to account for different mean firing rates. Then, for each grid cell pair, the spatial cross-correlation of these maps was obtained. The cross-correlations were smoothed using a Gaussian kernel with a standard deviation of 4 cm. Local maxima are identified by applying a threshold of 10% of the data range. For illustration, the local maximum that is closest to the center is identified, and the offset vector is defined as the vector connecting the center with the closest local maximum.

For calculation of the rotation of grid cell pairs between two trials *t*_1_ and *t*_2_, for each grid cell pair the cross-correlation of *t*_1_ is rotated repeatedly at multiples of 2° and the Pearson r correlation coefficient between the rotated cross-correlation of *t*_1_ and the crosscorrelation of *t*_1_ is obtained. The angle at which the mean correlation coefficient across grid cell pairs is maximized is defined as the rotation angle between the trials.

For determining the phase shift, for each grid cell pair, the cross-correlation of *t*_1_ is rotated to the rotation angle calculated previously and the cross-correlation between this rotated cross-correlation and the cross-correlation of *t*_2_ is obtained. The phase shift is defined as the maximum of this cross-correlation after averaging across grid cell pairs.

Based on the results of Supp. fig. 5, which are consistent with the literature ^73,74^, no module separation was performed prior to calculation of grid cell rotation and phase shift.

#### HD cell rotation, border cell rotation and directional firing score

To calculate the rotations of HD cells between two trials, the firing rate histogram per head direction was calculated for each trial, as described for the HD score calculation. The mean angle weighted by the firing rate was defined as the preferred head direction of the HD cell in that trial. To calculate the rotation, the difference of this preferred direction between the two trials was obtained.

For the rotation of border cells, the Cartesian position coordinates were first transformed to polar coordinates after centering at (0,0). The angle was then used to obtain a firing rate histogram (bin size = 10°, smoothing kernel = 10°). As for the HD cell rotation, the difference of the mean angles weighted by the firing rate of the two trials was taken as border cell rotation. In analogy to the HD score, the directional firing score was calculated using the mean vector length. To enhance the reliability of the border cell rotation only border cells with a directional firing score > 0.2 were included in the analysis.

#### LFP theta oscillations and running speed

For each channel, a frequency-power spectrogram was calculated using the *welch* function from the scipy package. The data were smoothed using a Gaussian kernel with a sigma of 1.5. The frequency with the highest peak within the theta range (4 - 12 Hz) was defined as the LFP theta frequency. The LFP theta power ratio was obtained from the ratio of the integrated signal within a 4 Hz frequency band around theta frequency and the sum of the integrated signal within two 2 Hz frequency bands above and below this interval. The channel with the highest theta power was used for the analysis.

For correlating theta power and theta frequency with the running speed, the running speed was calculated from the position data using the distance crossed between 2 data points and the sampling rate. The running speed was then smoothed with a Gaussian kernel with a sigma of 100 ms. Speed data that exceeded 60 cm/s was set to nan. Furthermore, the upper 10% of the speed data points were capped to avoid distortion of the linear approximation.

Theta power was calculated by applying a Butterworth bandpass filter for theta oscillations (4 - 12 Hz) to the LFP signal. The next step was to smooth the absolute signal with a Gaussian kernel with a sigma of 1250).

Theta frequency was obtained by identifying theta cycles in the LFP signal as described in the section on phase precession. The frequencies of the theta cycles are obtained from their durations and frequencies outside the theta range (4 - 12 Hz) are removed.

After interpolating the speed and the theta power or theta frequency data, a linear function was fitted to the correlation.

#### Phase precession and preferred theta phase

For the phase precession analysis, the phase in the theta cycle and the distance of the mouse from the entry point into the firing field for each spike were determined. The first step was to detect theta epochs on the channel with the highest waveform amplitude of the neuron. For this purpose, a Butterworth bandpass filter for theta (4 - 12 Hz) was applied to the LFP signal. Then, the absolute signal was convoluted with a Gaussian filter (sigma of 200 ms). The same analysis was performed for the delta range (2 - 4 Hz). Finally, theta epochs were defined as time periods of at least 500 ms for which the theta-delta ratio was greater than 2. For all theta epochs, theta cycles were identified by means of the positive-to-negative transitions of the signal. By obtaining the time point of each spike within its theta cycle, the phase was determined. The next step was to calculate the distance of the animal from the entry point into the firing field at the time the spike occurred. First, firing fields were defined as clusters of at least 10 contiguous bins in the firing rate map (bin size = 2 cm, smoothing kernel = 2 cm) for which the minimum peak rate was > 4 Hz and 1.5 times greater than the mean firing rate and the minimum firing rate for a cluster to be added to the field was 0.5 times the peak firing rate in the potential field. Passes were identified by the entry and exit points from the field and had to pass the following criteria (similar to the study by Schlesiger et al. ^26^): a minimal duration of 0.2 sec, a minimal mean speed of the animal of 3 cm/s and a minimum of 5 spikes. Lastly, the relative distance of the mouse at the time of the spike from the entry point into the fields compared to the length of the path (from the entry point to the exit point) was calculated. The regression slope and the corresponding p-value were derived from the circular-linear regression analysis developed by Kempter et al. ^75^. The regression analysis was either applied to all spikes fired during one path through a field or after pooling all spikes from all passes through that field, as specified by the figure legend.

The preferred theta phase of a neuron was determined by calculating the circular mean phase of all spikes. The mean vector length (MVL) of this preferred theta phase was used to quantify the degree of phase locking to the LFP theta rhythm.

#### Theta index

To determine the degree of theta-rhythmic firing, the theta index was calculated for each neuron. First, the instantaneous firing rate was obtained by binning the spike train into intervals of 10 ms and smoothing with a Gaussian kernel with a sigma of 10 ms. Next, the power spectrum of the instantaneous firing rate was calculated using the *welch* function from scipy and smoothed using a Gaussian kernel with a sigma of 1.5 bins. To avoid the necessity of obtaining a peak in theta range in the power spectrum, the theta index was calculated using the theta frequency of the LFP or 8 Hz if the latter was undefined. The theta index was defined as 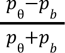 where *p*_θ_ and *p_b_* are the mean power in the 2 Hz frequency band around theta frequency and in the two 1 Hz frequency bands above and below, respectively.

To obtain a cutoff theta index to differentiate theta-rhythmic and non-theta-tuned HD cells, a kernel density estimation was applied to the distribution of theta indices from HD cells in young and mature control animals, using a Gaussian kernel with a bandwidth of 0.025. The local extrema were calculated, and the trough at 0.156 was used as a cutoff value.

#### Statistical analysis

For comparing dependent samples lacking a normal distribution, Mann-Whitney-Wilcoxon tests were used to compare the control group against the tau or amyloid mice. Two-sided Mann-Whitney-Wilcoxon tests were performed when impairments could result from a change in both directions (e.g. theta power), otherwise a one-sided version was employed. Due to multiple testing, the Holm-Bonferroni correction was applied. Results that turned out non-significant after this correction are highlighted in the figure legends. For circular-linear correlations, a Kempter regression analysis was performed. In the case of circular-circular correlations, the correlation coefficient and slope were obtained according to the work by Jammalamadaka ^76^.

For the behavioral experiments, unpaired t-tests were used to test for differences between two groups, while one-way ANOVA tests were employed for data with more than 2 groups. In case of detection of a significant difference, Tukey’s HSD test was used to calculate the p-values for the comparisons between the different groups. To test for interaction effects, a two-way ANOVA test was used.

## Supplementary figures

**Supplementary Fig. 1:**
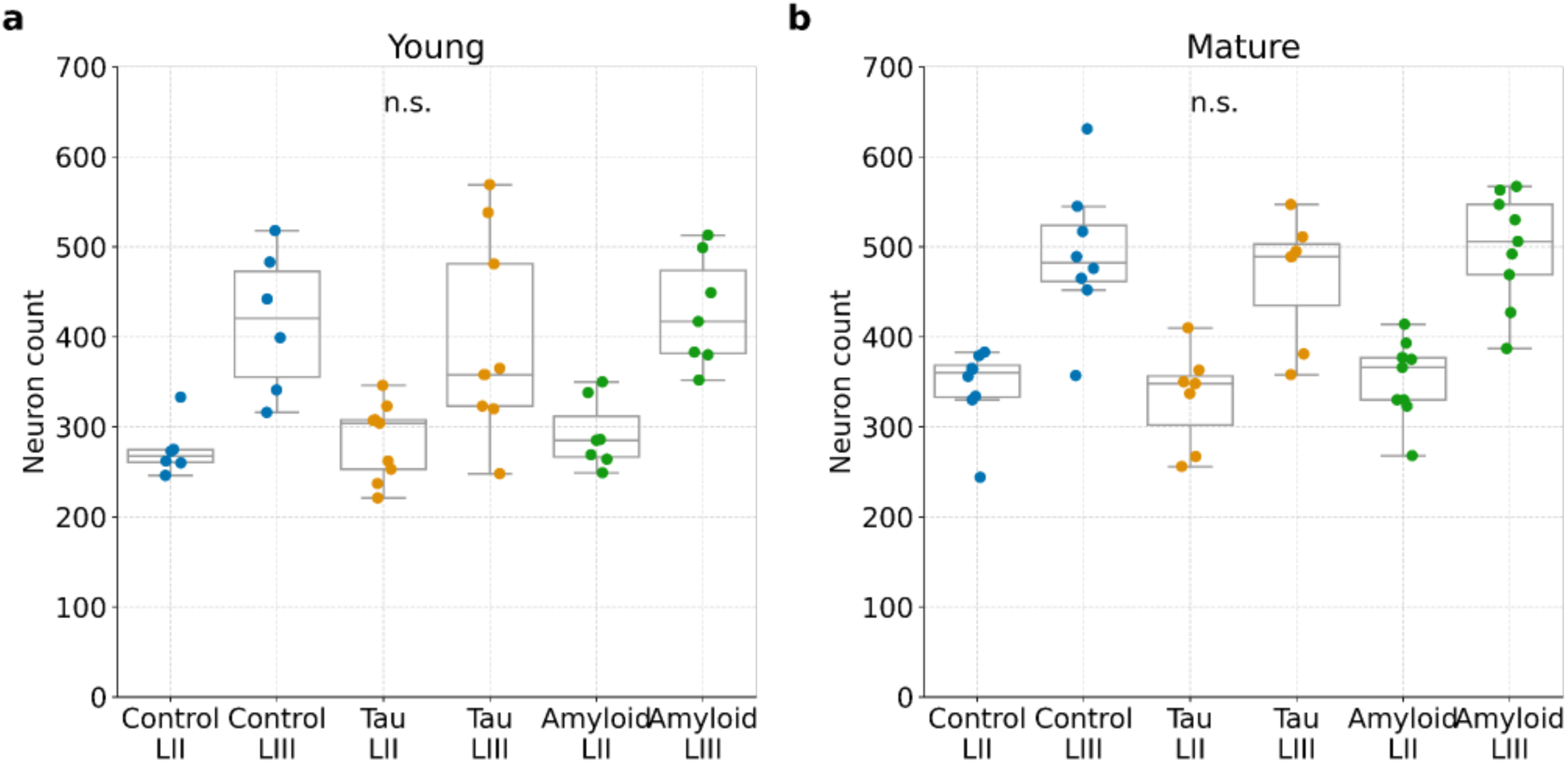
Preserved cell numbers in the superficial layers of MEC in young and mature tau and amyloid mice. (a) Neuron count in layer 2 (LII) and layer 3 (LIII) of MEC slices of young (5 months old) control (n = 6), tau (n = 9) and amyloid mice (n = 7). One-way ANOVA: LII: p = 0.737, LIII: p = 0.765 (b) Same as in a, but for mature (8 months old) control (n = 8), tau (n = 7) and amyloid mice (n = 9). One-way ANOVA: LII: p = 0.712, LIII: p = 0.662.

**Supplementary Fig. 2:**
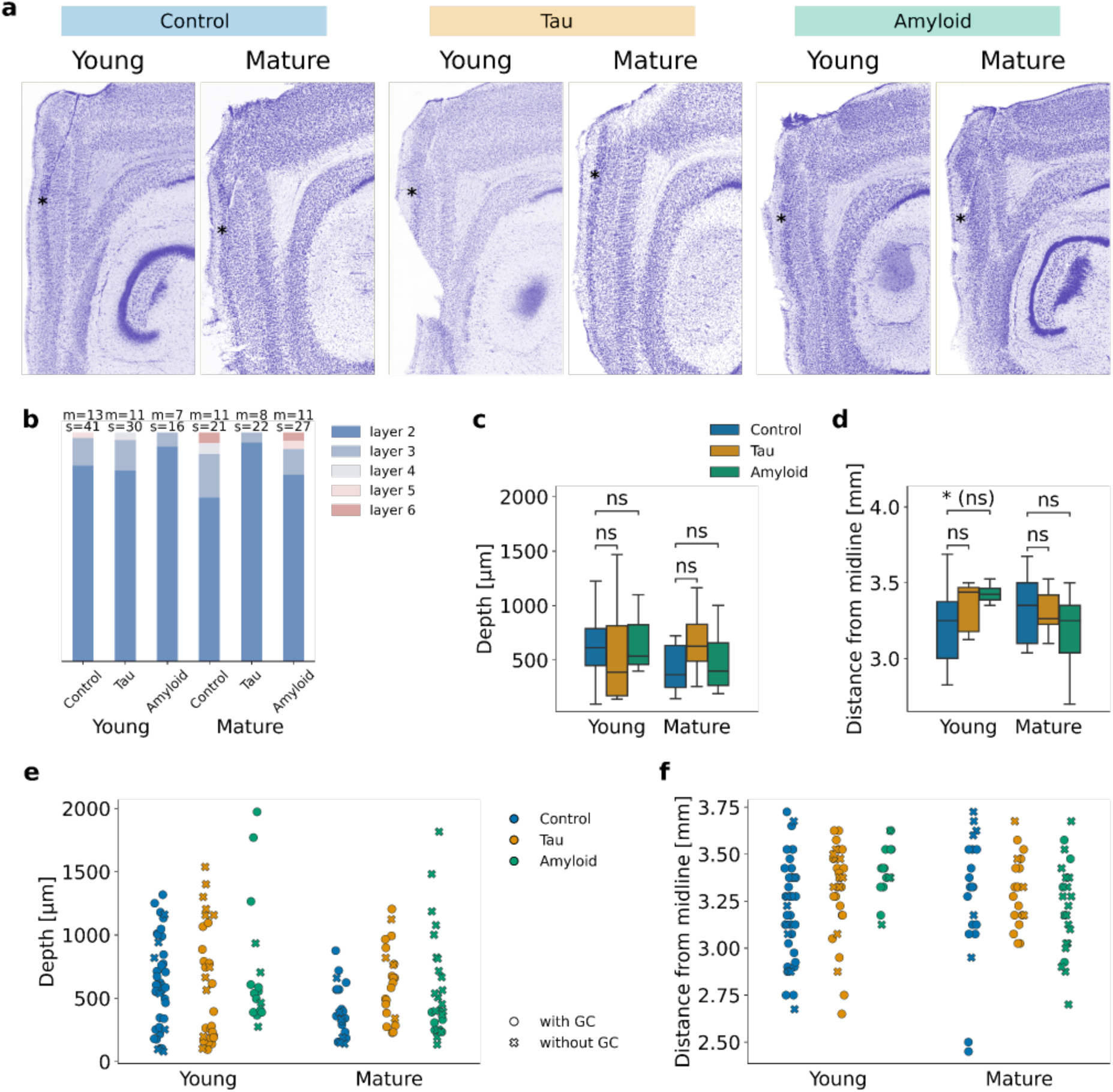
Homogeneous distribution of recording locations in control, tau and amyloid mice. (a) Brain sections with Nissl staining showing the location of one of the silicon probe shanks. Black asterisks highlight the tip of the shanks. (b) Distribution of shank tip locations across the layers in MEC. On top, the number of mice per group (m) and the number of shanks (s) is printed. Chi square test: df=5, p=0.296, ^2^ = 6.1. (c) Distribution of shank tip locations in the dorsoventral axis. The depth was defined as the distance in the dorsoventral axis between the tip of each shank and the dorsal border of the MEC. Mann-Whitney-Wilcoxon test, two-sided with Holm-Bonferroni correction, statistical unit: mouse: young control vs. young tau: p = 0.56, U = 75; young control vs. young amyloid: p = 0.82, U = 49; mature control vs. mature tau: p = 0.052, U = 20; mature control vs. mature amyloid: p = 0.43, U = 48 (d) Distribution of shank tip locations in the mediolateral axis (distance from midline). Mann-Whitney-Wilcoxon test, two-sided with Holm-Bonferroni correction, statistical unit: mouse: young control vs. young tau: p = 0.15, U = 41.5; young control vs. young amyloid: p = 0.026 (n.s. after Holm-Bonferroni correction), U = 17; mature control vs. mature tau: p = 1.0, U = 44; mature control vs. mature amyloid: p = 0.22, U = 79.5 (e) Same as (c), but each shank tip is shown. Shanks on which no grid cells were recorded are marked by a cross. The legends apply to (e) and (f). (f) Same as (d), but each shank tip is shown. Shanks on which no grid cells were recorded are marked by a cross.

**Supplementary Fig. 3:**
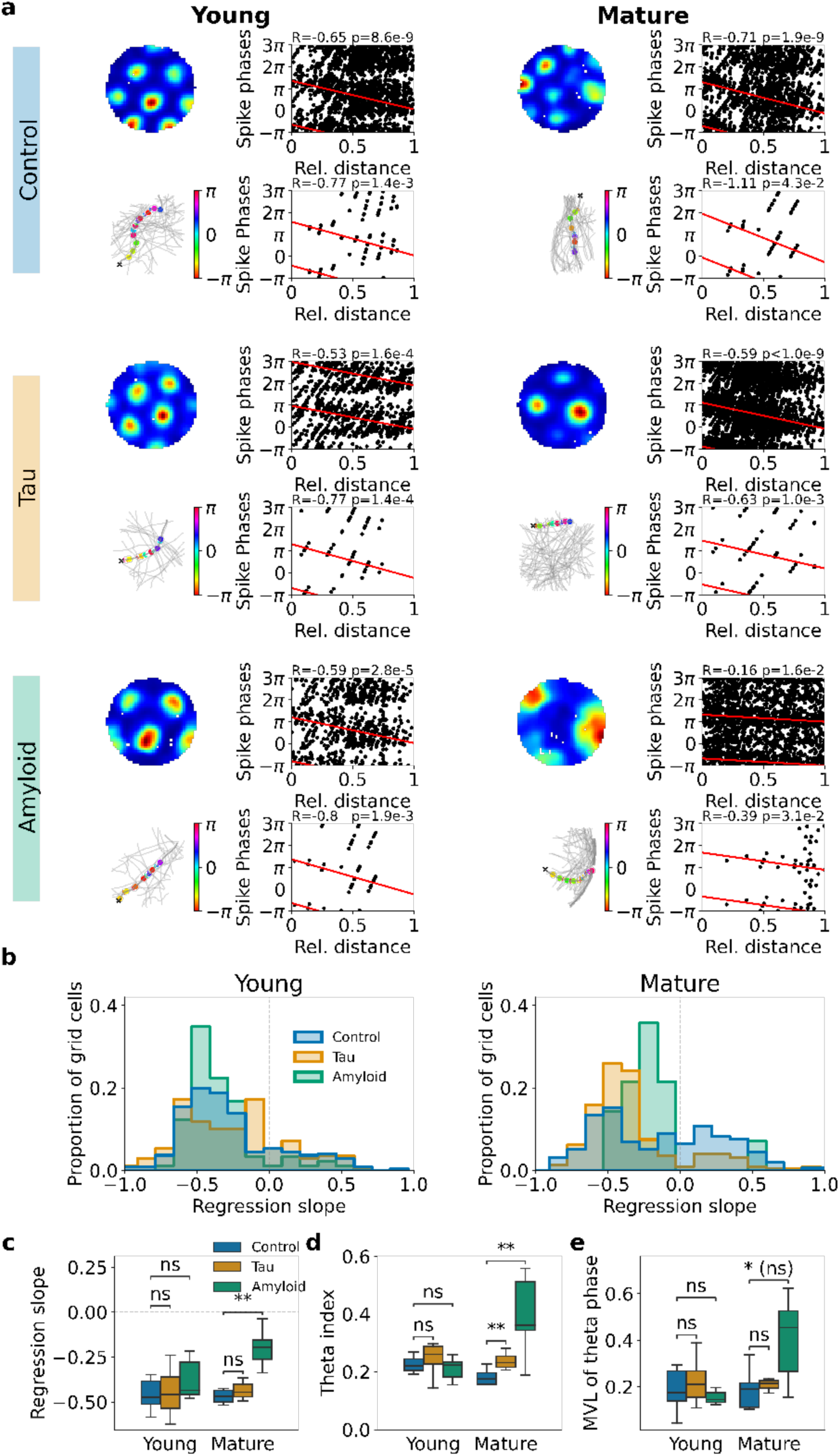
Reduced phase precession slope and enhanced theta modulation in grid cells in mature amyloid mice. (a) For each of the 6 groups an example grid cell featuring phase precession is presented. The top left diagram shows the firing rate map, and the top right diagram depicts the correlation between the spike phases and the relative distance of the mouse from the entry point into the field. For better visualization of the circular data, 2 theta cycles are shown. The plot refers to the field with the highest peak firing rate. The regression slope (R) and the corresponding p-value (p) derived from the Kempter circular-linear regression analysis are printed above the plot. The bottom left diagram shows the analyzed paths through the field (light gray); an example path is marked in dark gray. The spikes with color-coded phases are plotted according to their position on the path. The bottom right diagram represents the correlation between these spike phases and the relative distance of the mouse from the entry point into the field. The regression slope (R) and the p-value (p) are printed above. (b) The histograms of the regression slopes are shown for the young and mature cohorts separately. The spikes were pooled by field for the calculation of the regression slopes. The slopes were then averaged across all fields per grid cell. Wilcoxon signed-rank test, two-sided with Holm-Bonferroni correction, statistical unit: cell: young control: n = 245, p = 1.8*10^-22^, U = 4240; young tau: n = 168, p = 2.5*10^-14^, U = 2287; young amyloid: n = 89, p = 3.9*10^-13^, U = 228; mature control: n = 157, p = 2.6*10^-5^, U = 3801; mature tau: n = 227, p = 5.2*10^-26^, U = 2490; mature amyloid: n = 14, p = 0.013, U = 14 (c) Boxplots showing the distribution of regression slopes of phase-precessing grid cells (at least one firing field with a significant negative regression slope) across cohorts. Mann-Whitney-Wilcoxon test, two-sided with Holm-Bonferroni correction, statistical unit: mouse: young control vs. young tau: p = 0.94, U = 48; young control vs. young amyloid: p = 0.30, U = 22; mature control vs. mature tau: p = 0.62, U = 20; mature control vs. mature amyloid: p = 0.0051, U = 1 (d) Theta modulation of grid cells as quantified by the theta index. The theta index is calculated from the instantaneous firing rate power spectrum and is defined as the ratio of the theta power close to the peak theta frequency f∓1Hz and the power at baseline (between f-3Hz and f-2Hz and between f+2Hz and f+3Hz). Mann-Whitney-Wilcoxon test, two-sided with Holm-Bonferroni correction, statistical unit: mouse: young control vs. young tau: p = 0.10, U = 38; young control vs. young amyloid: p = 0.76, U = 50; mature control vs. mature tau: p = 0.0052, U = 6; mature control vs. mature amyloid: p = 0.007, U = 3 (e) Locking of grid cells to LFP theta as quantified by the mean vector length of the spike phases. Mann-Whitney-Wilcoxon test, two-sided with Holm-Bonferroni correction, statistical unit: mouse: young control vs. young tau: p = 0.73, U = 59; young control vs. young amyloid: p = 0.39, U = 57; mature control vs. mature tau: p = 0.21, U = 19; mature control vs. mature amyloid: p = 0.042 (n.s. after Holm-Bonferroni correction), U = 7

**Supplementary Fig. 4:**
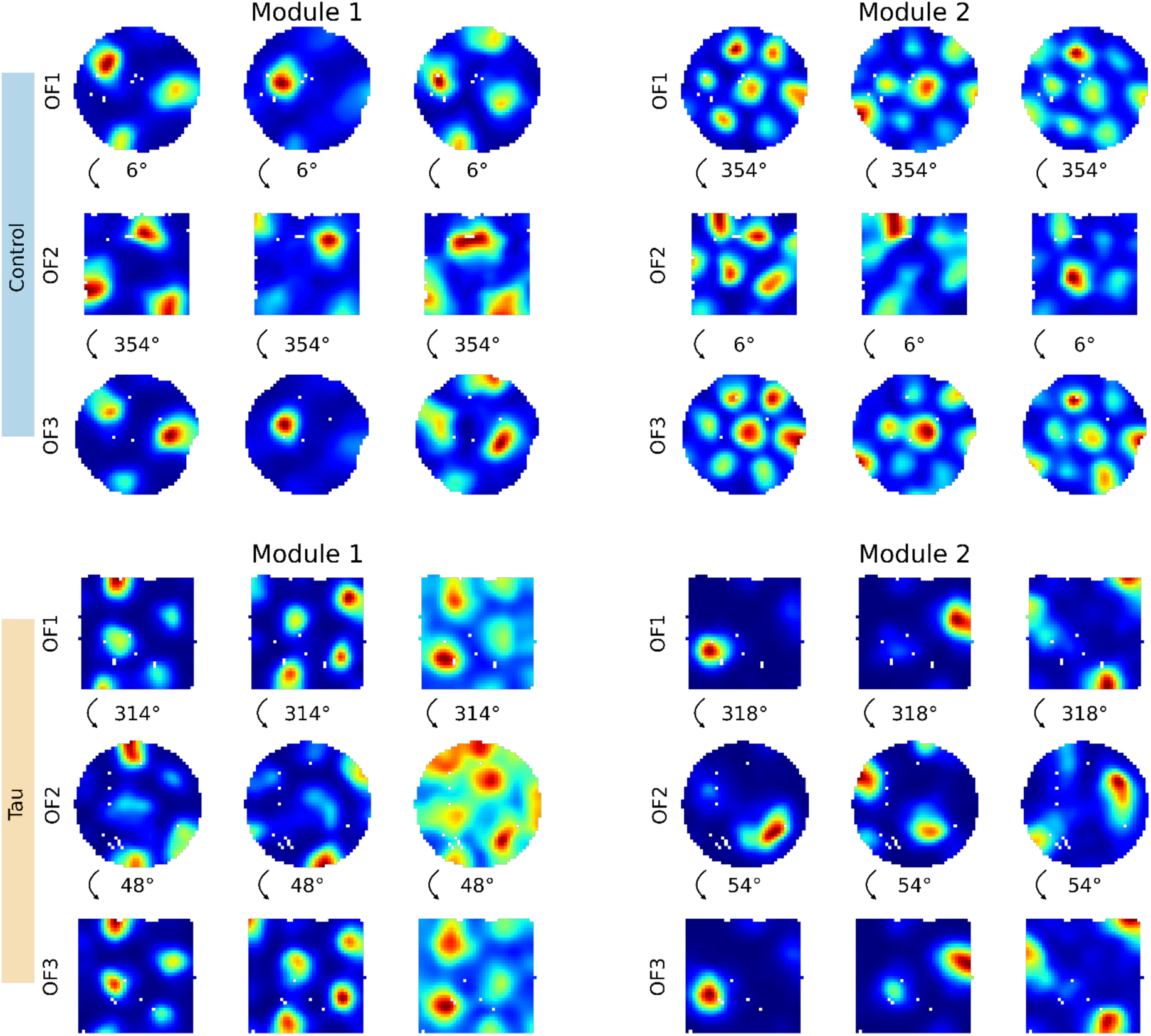
Coherent inter-trial rotations in recordings with two grid modules. Two example sessions with 2 different grid modules, one from a young control mouse (above), one from a young tau mouse (below). The module with the higher number of grid cells is shown on the left, the other one on the right. For each module, the firing rate maps for the 3 trials (open field (OF) 1-3 of 3 example grid cells are shown. The rotation of the grid pattern (in counter-clockwise direction) is printed between the trials. Recordings of 2 different modules with at least 2 grid cells each were rare (young control: 3 sessions, young tau: 2 sessions, young amyloid: 0 sessions, mature control: 2 sessions, mature tau: 5 sessions, mature amyloid: 0 sessions).

**Supplementary Fig. 5:**
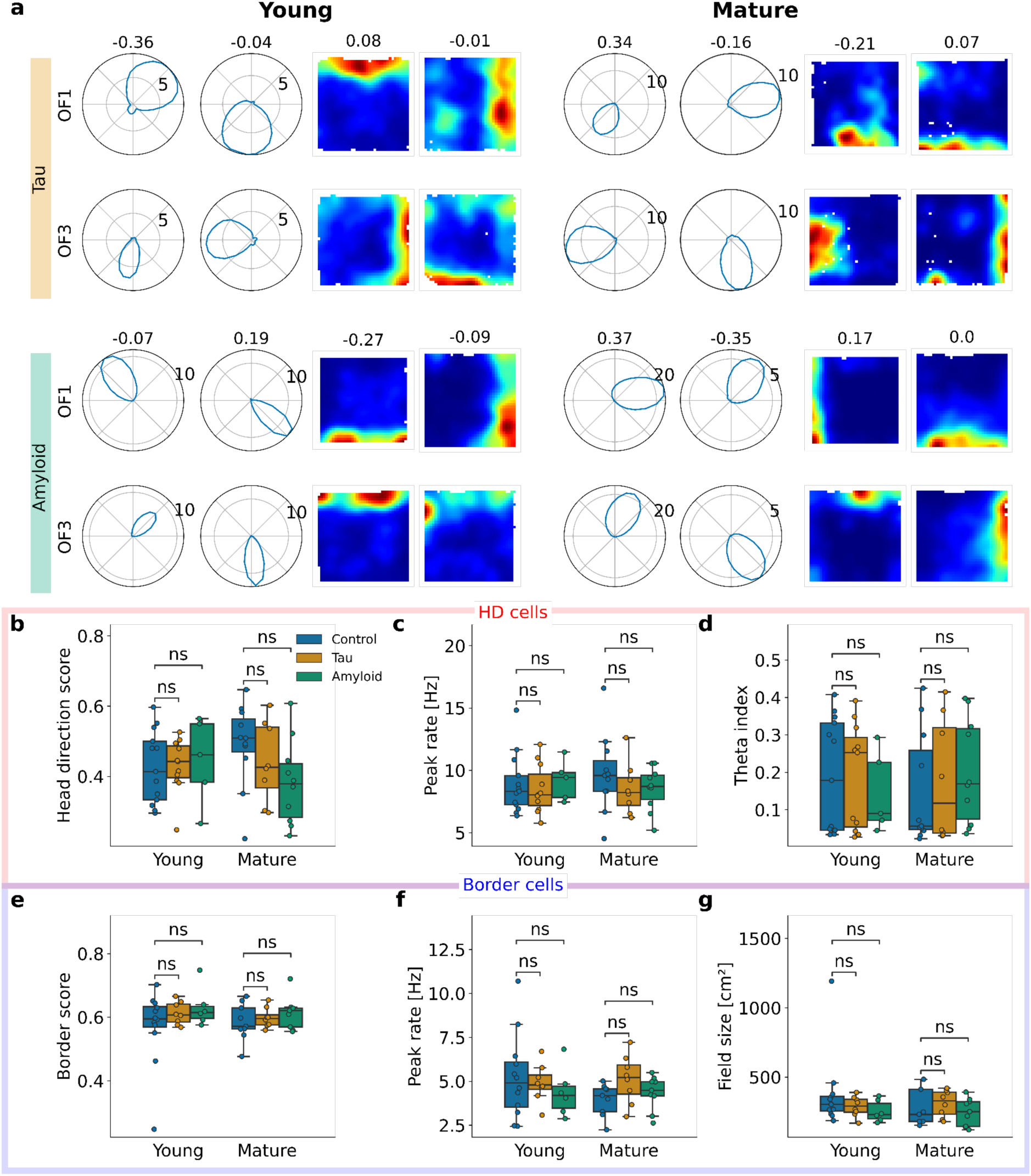
Basic properties of head-direction cells and border cells in young and mature AD mice are preserved. (a) Further examples of HD cells and border cells that display an inter-trial instability. Two examples for HD cells (polar HD histograms) and border cells (firing rate maps) in the first and last open field trial (OF1 and OF3) for young and mature tau and amyloid mice are shown. The Pearson correlation coefficient between firing rate maps or HD rate histograms is shown above map pairs. (b-d) Boxplots showing the HD score, peak firing rate and theta index of HD cells. Each data point is the median of the HD cells recorded in one mouse. The legend applies to b-g. (b) Head direction score of HD cells. Mann-Whitney-Wilcoxon test, two-sided with Holm-Bonferroni correction, statistical unit: mouse: young control vs. young tau: p = 0.91, U = 69; young control vs. young amyloid: p = 0.58, U = 30; mature control vs. mature tau: p = 0.40, U = 55; mature control vs. mature amyloid: p = 0.073, U = 81 (c) Peak firing rate of HD cells. Mann-Whitney-Wilcoxon test, two-sided with Holm-Bonferroni correction, statistical unit: mouse: young control vs. young tau: p = 0.77, U = 77; young control vs. young amyloid: p = 0.50, U = 25; mature control vs. mature tau: p = 0.31, U = 57; mature control vs. mature amyloid: p = 0.42, U = 67 (d) Theta index of HD cells. Mann-Whitney-Wilcoxon test, two-sided with Holm-Bonferroni correction, statistical unit: mouse: young control vs. young tau: p = 0.91, U = 74; young control vs. young amyloid: p = 0.85, U = 35; mature control vs. mature tau: p = 0.97, U = 45; mature control vs. mature amyloid: p = 0.31, U = 40 (e-g) Boxplots showing the border score, peak firing rate and field size of border cells. Each data point is the mean of the border cells in one mouse. (e) Border score of border cells. Mann-Whitney-Wilcoxon test, two-sided with Holm-Bonferroni correction, statistical unit: mouse: young control vs. young tau: p = 0.57, U = 40; young control vs. young amyloid: p = 0.55, U = 29; mature control vs. mature tau: p = 0.48, U = 28; mature control vs. mature amyloid: p = 0.54, U = 33 (f) Peak firing rate of border cells. Mann-Whitney-Wilcoxon test, two-sided with Holm-Bonferroni correction, statistical unit: mouse: young control vs. young tau: p = 0.97, U = 47; young control vs. young amyloid: p = 0.93, U = 18; mature control vs. mature tau: p = 0.093, U = 18; mature control vs. mature amyloid: p = 0.55, U = 43 (g) Mean field size of border cells. Mann-Whitney-Wilcoxon test, two-sided with Holm-Bonferroni correction, statistical unit: mouse: young control vs. young tau: p = 0.68, U = 54; young control vs. young amyloid: p = 0.21, U = 50; mature control vs. mature tau: p = 0.60, U = 30; mature control vs. mature amyloid: p = 0.60, U = 47

**Supplementary Fig. 6:**
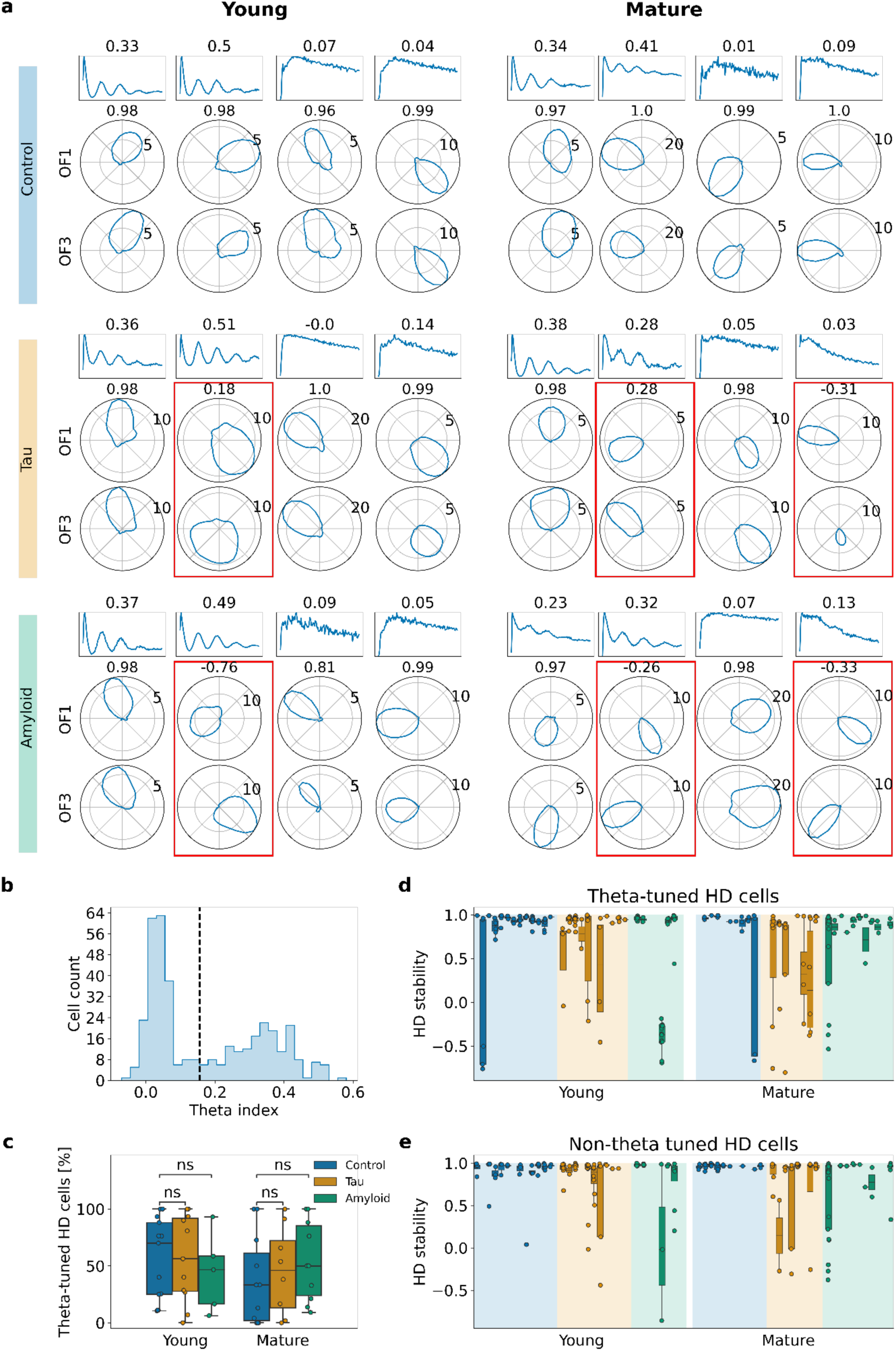
Theta-tuned and non-theta-tuned head direction cells exhibit inter-trial instability in mice of both genotypes. (a) Examples for theta-tuned and non-theta-tuned HD cells for each cohort. Top row: Normalized spike-time autocorrelogram from 0 to 0.5 sec. The theta index is printed above. Second and third row: Polar HD histograms of the first and last open field trial (OF1 and OF3). The Pearson correlation coefficient between firing rate maps or HD rate histograms is shown above map pairs. Red boxes mark examples with a correlation coefficient < 0.5. (b) Histogram of theta indices in young and mature control mice. The dashed black line at x = 0.156 marks the limit between theta-tuned and non-theta-tuned HD cells based on a Gaussian kernel density analysis. (c) Proportions of theta-tuned HD cells with respect to all HD cells. Each dot represents a mouse. Mann-Whitney-Wilcoxon test, two-sided with Holm-Bonferroni correction, statistical unit: mouse: young control vs. young tau: p = 0.84, U = 67.5; young control vs. young amyloid: p = 0.49, U = 40; mature control vs. mature tau: p = 0.56, U = 36.5; mature control vs. mature amyloid: p = 0.24, U = 38 (d) HD stability of theta-tuned HD cells per mouse. Each bar represents the data from one mouse. Each dot is the median Pearson correlation coefficient of theta-tuned HD cells co-recorded in one session. Mann-Whitney-Wilcoxon test, one-sided with Holm-Bonferroni correction, statistical unit: session: young control vs. young tau: p = 0.13, U = 1239; young control vs. young amyloid: p = 0.011, U = 1018; mature control vs. mature tau: p = 0.016, U = 3550; mature control vs. mature amyloid: p = 0.059, U = 3180 (e) Same as (d), but for non-theta-tuned HD cells. Mann-Whitney-Wilcoxon test, one-sided with Holm-Bonferroni correction, statistical unit: session: young control vs. young tau: p = 0.045 (n.s. after Holm-Bonferroni correction), U = 1093; young control vs. young amyloid: p = 0.28, U = 61; mature control vs. mature tau: p = 0.0054, U = 71.9; mature control vs. mature amyloid: p = 0.018 (n.s. after Holm-Bonferroni correction), U = 77.8

**Supplementary Fig. 7:**
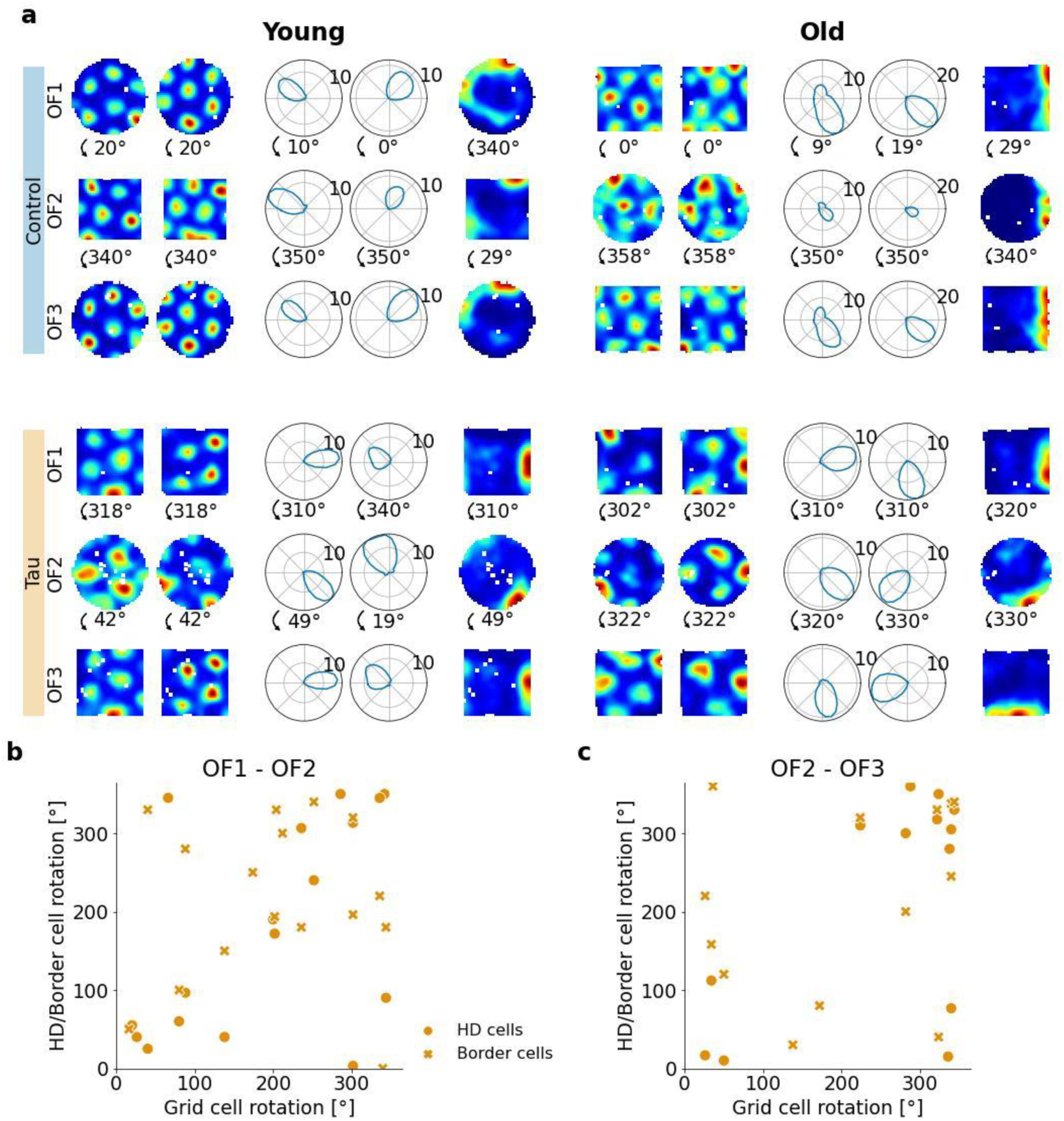
Rotations in grid cells, HD cells and border cells are correlated in tau mice. (a) Example sessions with 2 grid cells, 2 HD cells and 1 border cell recorded simultaneously for young and mature control and tau mice. The firing rate maps (grid cells, border cell) or polar HD histograms (HD cells) for each open field trial (OF1, OF2, OF3) are shown. The rotation of the cells (in counter-clockwise direction) is printed between the trials. (b) Scatter plot showing the correlation between the grid cell rotation and the rotation of HD or border cells between the first and second open field trials in mature tau mice. Each data point is a session with at least 2 co-recorded grid cells rotating at least 15° in either direction and at least 1 HD (n=17 sessions) or border cell (n=16 sessions) recorded simultaneously. Circular correlation: rho = 0.564, p = 0.0018 (c) Same as (b), but for rotations between the second and last open field trial. 13 sessions for HD cells and 14 sessions for border cells. Circular correlation: rho = 0.527, p = 0.0065

**Supplementary Fig. 8:**
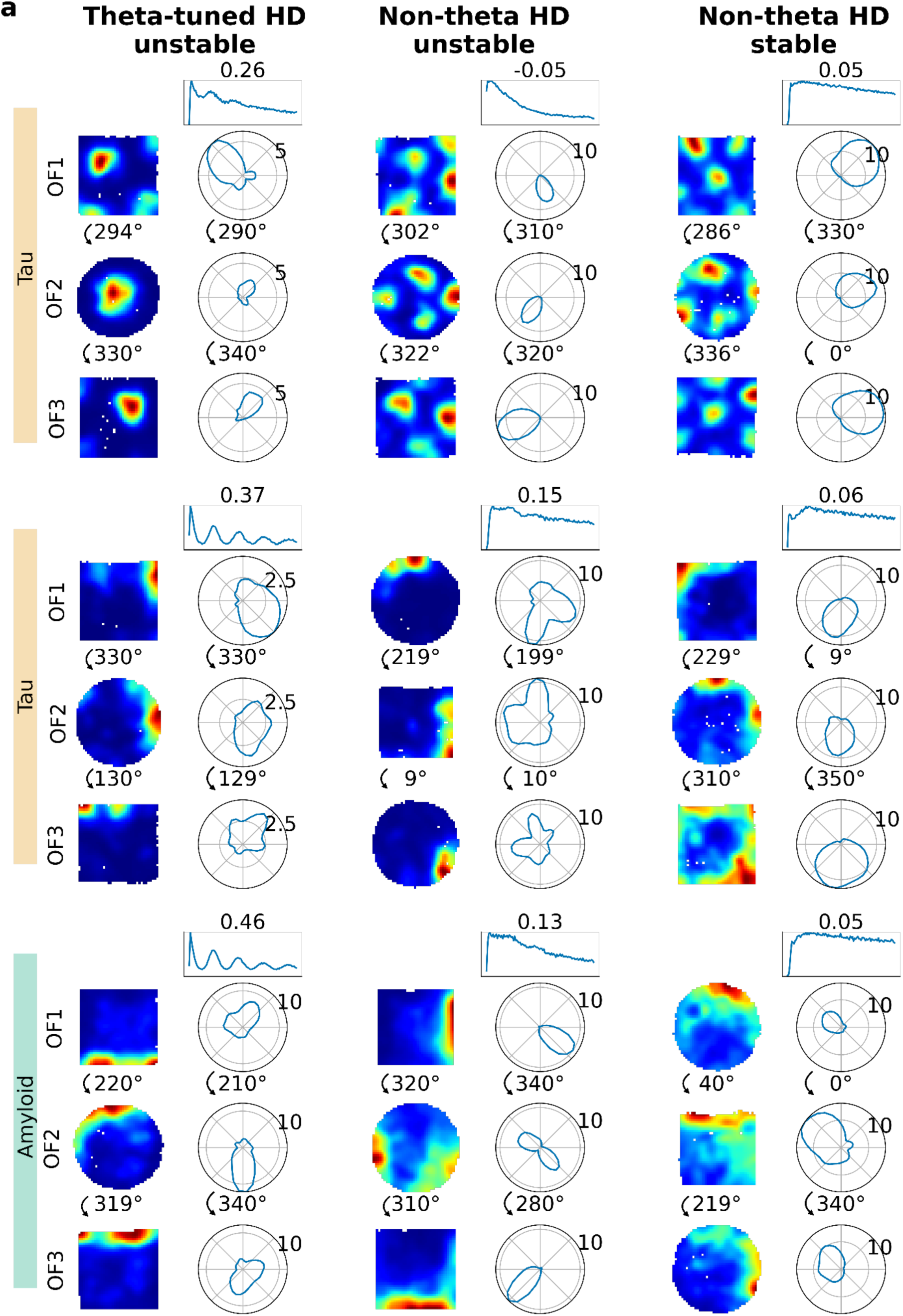
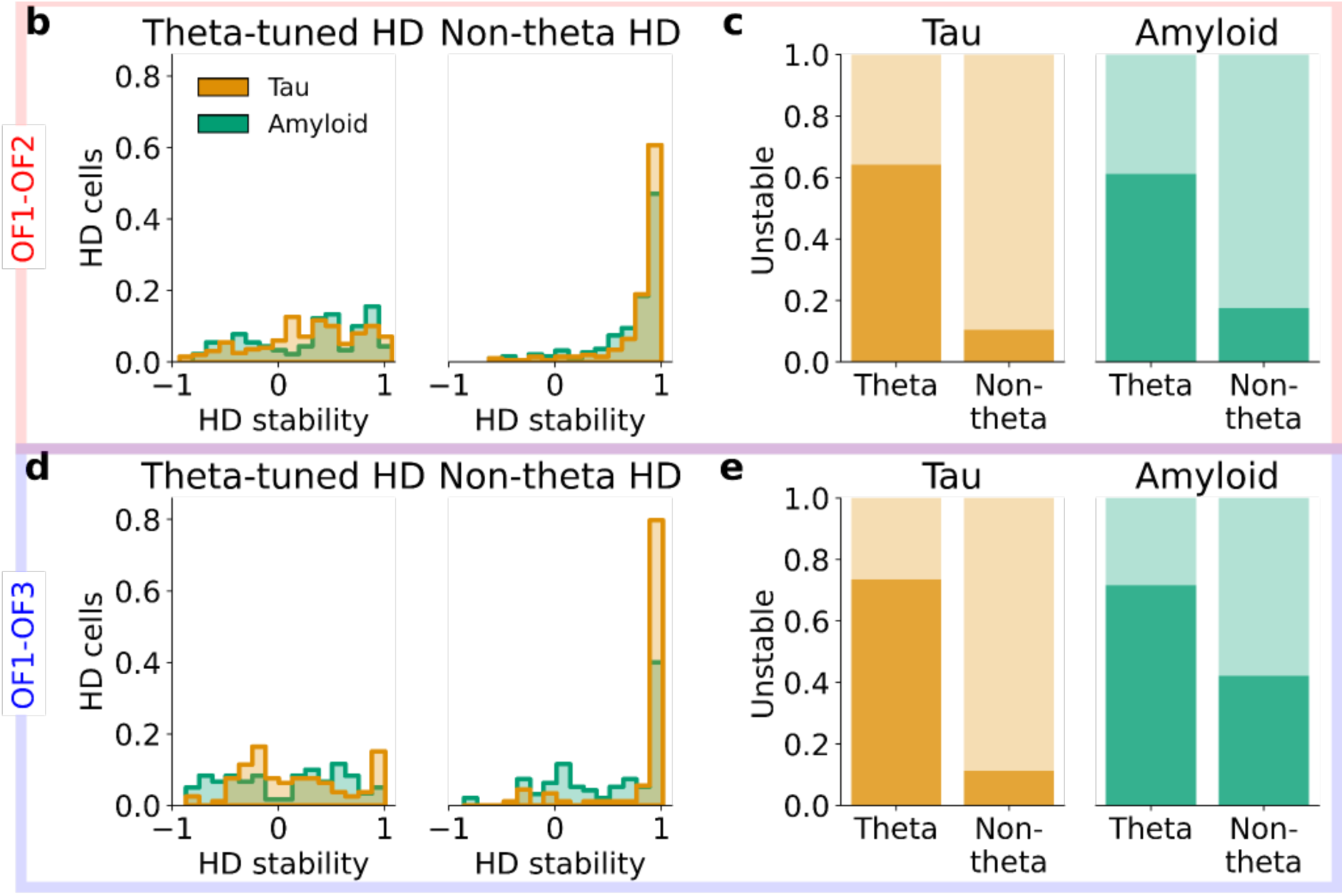
Theta-tuned HD cells consistently show inter-trial instability in sessions with unstable grid or border cells, whereas non-theta-tuned HD cells remain mostly stable in tau and amyloid mice. (a) Top: Example sessions with unstable theta-tuned HD cells (left), unstable non-theta-tuned HD cells (middle) and stable non-theta-tuned HD cells (right) in sessions with unstable grid cells in tau mice (top row) or border cells in tau mice (second row) or border cells in amyloid mice (third row). The firing rate maps (grid cells, border cell) or polar HD histograms (HD cells) for each open field trial (OF1, OF2, OF3) are shown. The rotation of the cells (in counter-clockwise direction) is printed between the trials. The normalized spike-time autocorrelogram from 0 to 0.5 sec for the HD cell is plotted above the polar HD histograms with the theta index printed above. (b) Histogram of HD stability between the first and second open field trial in theta-tuned HD cells (left) and non-theta-tuned HD cells (right) for tau and amyloid mice. ^(c)^ Proportion of unstable theta-tuned HD cells and non-theta-tuned HD cells in tau (left) and amyloid mice (right) in sessions with unstable grid cells and/or border cells. Stability was defined as a map cross-correlation or an HD cross-correlation > 0.6. Chi-square test: Tau: ^2^ = 120.6, p < 10^-9^, amyloid: ^2^ = 51.8, p < 10^-9^ (d) Same as b, but for the first and last open field trials. (e) Same as c, but for the first and last open field trials. Chi-square test: Tau: ^2^ = 64.6, p < 10^-9^, amyloid: ^2^ = 11.8, p = 0.0006

**Supplementary Fig. 9:**
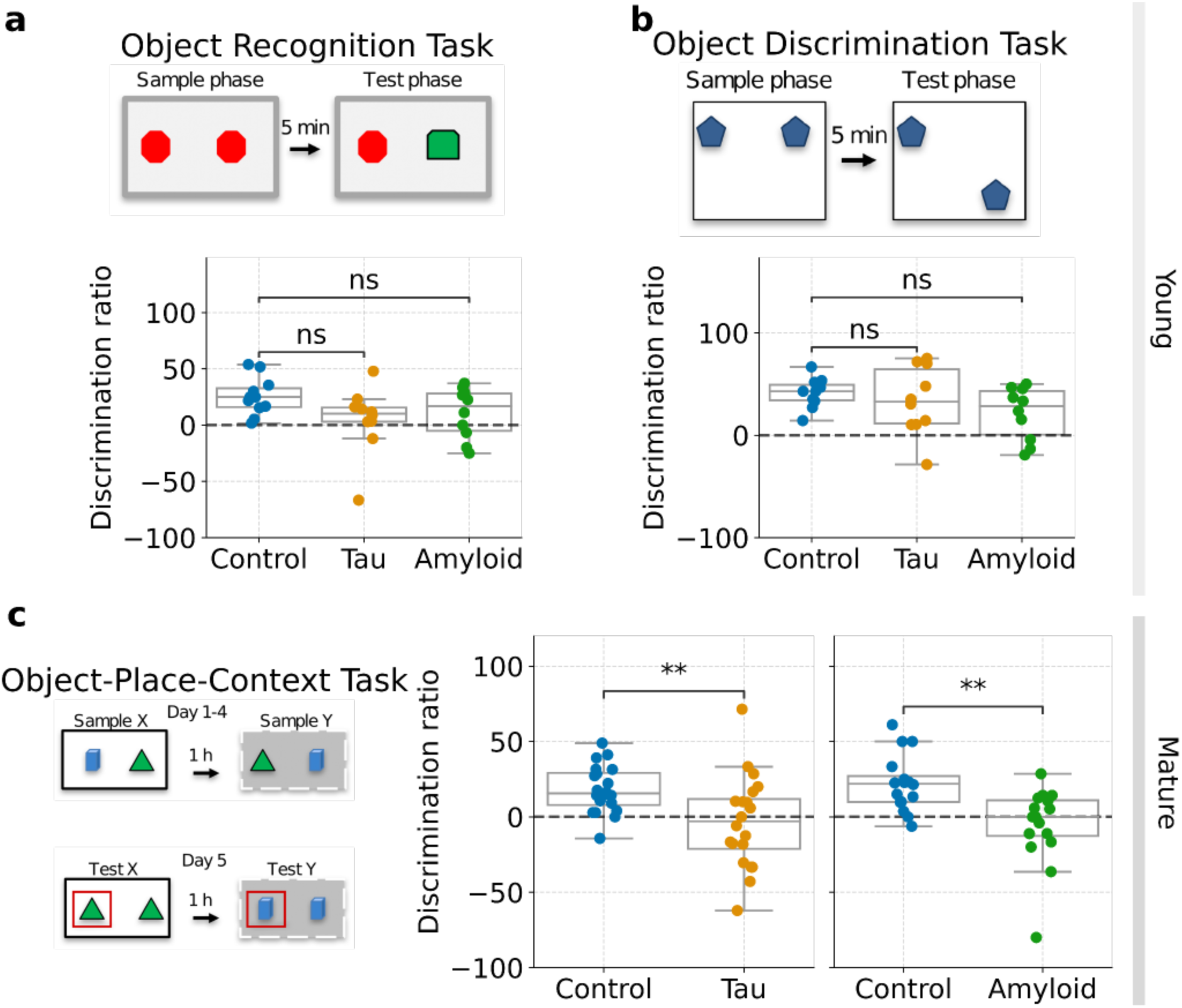
Intact object recognition and discrimination and impaired object-place-context coding in young and in mature tau and amyloid mice respectively. (a + b) Object recognition task (OR, a) and object displacement task (OD, b) were used to assess short-term recognition memory in young control, tau and amyloid mice. The discrimination ratio is calculated as follows: (L2 - L1)/(L1+L2)*100, where L1 and L2 are the exploration times for object 1 and object 2, respectively. Mice naturally tend to spend more time exploring the novel object or the object in the novel location. (a) Top: Experimental protocol of the OR task: In the test phase, one of the familiar objects (red) was replaced by a novel object (green). Bottom: Discrimination ratios for young control (n = 11), tau mice (n = 10) and amyloid mice (n = 10). one-way ANOVA: p = 0.1275, F(2,28) = 2.219. (b) Top: Experimental protocol of the OD task: In the test phase, one of the objects is moved to a different location. Bottom: Discrimination ratios for young control (n = 11), tau mice (n = 10) and amyloid mice (n = 10). one-way ANOVA: p = 0.2037, F(2,28) = 1.685. (c) Left column: Protocol of the Object-place-context task (OPC): The mice underwent 2 sample sessions for 4 consecutive days and two test sessions on day 5 (one in context X and one in context Y). The novel configurations in the test trials are highlighted by red frames. Middle Column: Boxplots indicating the discrimination ratios for mature control and tau mice (n = 10 mice per group, 2 values per mouse). The discrimination ratio is calculated as follows: (L2 - L1)/(L1+L2)*100, where L1 and L2 are the exploration times for object 1 and object 2, respectively. Mice naturally tend to spend more time exploring the new object-place-context configurations. Chance performance = 0%. Unpaired t-test: mature control vs. tau p = 0.0081. Right column: Same for mature control and amyloid mice (n = 8 mice per group, 2 values per mouse). Unpaired t-test: mature control vs. amyloid p = 0.0015.

**Supplementary table 1.1:** Numbers of sessions, cells, grid cells, head-direction cells, border cells and fast-spiking cells per mouse for young control mice

| Mouse | Sessions | Cells | Grid Cells | HD Cells | Border Cells | FS Cells |
| --- | --- | --- | --- | --- | --- | --- |
| bt068 | 11 | 383 | 104 | 9 | 3 | 40 |
| bt072 | 10 | 98 | 13 | 7 | 1 | 9 |
| bt0975 | 15 | 238 | 46 | 19 | 1 | 41 |
| bt1021 | 8 | 382 | 44 | 10 | 35 | 29 |
| bt1296 | 7 | 272 | 56 | 42 | 12 | 23 |
| bt1316 | 7 | 190 | 5 | 5 | 5 | 12 |
| bt1617 | 10 | 209 | 20 | 5 | 4 | 33 |
| bt3018 | 5 | 85 | 8 | 21 | 10 | 6 |
| bt3019 | 4 | 68 | 1 | 15 | 1 | 9 |
| bt3198 | 12 | 708 | 8 | 38 | 44 | 51 |
| bt3565 | 15 | 159 | 13 | 18 | 4 | 23 |
| bt6544 | 24 | 508 | 35 | 55 | 19 | 52 |
| bt8564 | 5 | 87 | 27 | 4 | 0 | 14 |

**Supplementary table 1.2:** Numbers of sessions, cells, grid cells, head-direction cells, border cells and fast-spiking cells per mouse for young tau mice

| Mouse | Sessions | Cells | Grid Cells | HD Cells | Border Cells | FS Cells |
| --- | --- | --- | --- | --- | --- | --- |
| bt0920 | 10 | 131 | 2 | 43 | 6 | 27 |
| bt1003 | 7 | 169 | 8 | 31 | 8 | 22 |
| bt1009 | 13 | 434 | 135 | 52 | 14 | 70 |
| bt1010 | 3 | 112 | 2 | 6 | 4 | 14 |
| bt1018 | 19 | 503 | 20 | 41 | 14 | 82 |
| bt1086 | 13 | 200 | 2 | 11 | 3 | 35 |
| bt1093 | 7 | 203 | 31 | 16 | 4 | 18 |
| bt1301 | 4 | 133 | 26 | 7 | 1 | 29 |
| bt1335 | 11 | 101 | 0 | 5 | 0 | 16 |
| bt1491 | 6 | 98 | 8 | 20 | 0 | 6 |
| bt1516 | 4 | 14 | 1 | 3 | 0 | 1 |

**Supplementary table 1.3:** Numbers of sessions, cells, grid cells, head-direction cells, border cells and fast-spiking cells per mouse for young amyloid mice

| Mouse | Sessions | Cells | Grid Cells | HD Cells | Border Cells | FS Cells |
| --- | --- | --- | --- | --- | --- | --- |
| bt0378 | 11 | 212 | 46 | 29 | 8 | 28 |
| bt0390 | 12 | 592 | 2 | 79 | 19 | 71 |
| bt0391 | 5 | 65 | 27 | 0 | 0 | 9 |
| bt0399 | 4 | 39 | 6 | 0 | 2 | 1 |
| bt0400 | 12 | 396 | 1 | 58 | 19 | 31 |
| bt0484 | 6 | 307 | 25 | 45 | 3 | 27 |
| bt0496 | 12 | 815 | 7 | 127 | 21 | 105 |

**Supplementary table 1.4:** Numbers of sessions, cells, grid cells, head-direction cells, border cells and fast-spiking cells per mouse for mature control mice

| Mouse | Sessions | Cells | Grid Cells | HD Cells | Border Cells | FS Cells |
| --- | --- | --- | --- | --- | --- | --- |
| bt0502 | 5 | 44 | 1 | 20 | 1 | 10 |
| bt1063 | 9 | 359 | 90 | 23 | 12 | 30 |
| bt1133 | 9 | 225 | 16 | 25 | 4 | 25 |
| bt1188 | 11 | 180 | 2 | 24 | 5 | 34 |
| bt1197 | 5 | 47 | 2 | 4 | 2 | 7 |
| bt1238 | 2 | 114 | 0 | 4 | 1 | 4 |
| bt1258 | 6 | 49 | 17 | 2 | 0 | 7 |
| bt4232 | 7 | 136 | 48 | 12 | 5 | 14 |
| bt64245 | 8 | 225 | 57 | 4 | 0 | 26 |
| bt64246 | 8 | 95 | 22 | 11 | 1 | 24 |
| bt9320 | 3 | 35 | 0 | 6 | 1 | 2 |

**Supplementary table 1.5:** Numbers of sessions, cells, grid cells, head-direction cells, border cells and fast-spiking cells per mouse for mature tau mice

| Mouse | Sessions | Cells | Grid Cells | HD Cells | Border Cells | FS Cells |
| --- | --- | --- | --- | --- | --- | --- |
| bt0962 | 18 | 441 | 118 | 26 | 7 | 68 |
| bt0963 | 7 | 227 | 6 | 23 | 16 | 20 |
| bt1047 | 12 | 639 | 139 | 34 | 21 | 50 |
| bt1056 | 8 | 229 | 34 | 16 | 8 | 33 |
| bt1070 | 12 | 671 | 9 | 106 | 27 | 136 |
| bt1181 | 7 | 161 | 34 | 11 | 4 | 17 |
| bt1221 | 7 | 397 | 31 | 76 | 11 | 40 |
| bt1223 | 9 | 568 | 0 | 48 | 7 | 76 |

**Supplementary table 1.6:** Numbers of sessions, cells, grid cells, head-direction cells, border cells and fast-spiking cells per mouse for mature amyloid mice

| Mouse | Sessions | Cells | Grid Cells | HD Cells | Border Cells | FS Cells |
| --- | --- | --- | --- | --- | --- | --- |
| bt0337 | 29 | 768 | 0 | 67 | 18 | 161 |
| bt0338 | 9 | 296 | 0 | 2 | 2 | 21 |
| bt0372 | 5 | 154 | 0 | 11 | 6 | 15 |
| bt0379 | 3 | 53 | 10 | 2 | 0 | 7 |
| bt0380 | 6 | 299 | 2 | 33 | 8 | 24 |
| bt0383 | 3 | 61 | 1 | 21 | 2 | 7 |
| bt0384 | 2 | 144 | 1 | 34 | 4 | 14 |
| bt0410 | 13 | 71 | 4 | 4 | 2 | 8 |
| bt0504 | 8 | 47 | 0 | 2 | 2 | 3 |
| bt562 | 9 | 15 | 0 | 0 | 0 | 2 |
| bt565 | 12 | 196 | 0 | 29 | 11 | 12 |

## Acknowledgements

We thank the imaging core facility and the animal facility of the DKFZ for their technical assistance. We thank S. Desikan for her help with immunohistochemical stainings. We thank P. Klein for his help with experiments and analysis. We thank R. Groeneveld and T. Fabian for their help with experiments during their internships in the lab. We thank A. Caputi and M. Schlesiger for the insightful discussions and inspiring feedback on the manuscript.

This work was made possible by the Deutsche Forschungsgemeinschaft via Individual Research Grants (the SFB-1436/2 Project-ID 425899996, the SFB/TRR384 IN-CODE, the Advanced ERC grant Project-ID 101142587 and a NRRP grant, project MNESYS/PE0000006/DN.1553 11.10.2022 to H.M.). We thank the Lautenschläger Foundation for supporting the Department of Clinical Neurobiology and the Neurology Clinic (H.M. and W.W.).

## Author contributions

Conceptualization: HM

Methodology and experiments: BT, EF, J-JP, HO

Data analysis: BT, EF

Funding acquisition: HM, WW

Writing -original draft: BT, EF

Writing -review and editing: BT and HM with the help of the other auhors

## Data and materials availability

The spike trains, animal position data, and task event logs will be published on the open-access Dryad platform. The computer code for data analysis will be accessible on a public GitHub repository.

